# Extreme drought disrupts the bottom-up regulation of ground-dwelling invertebrate diversity under climate warming

**DOI:** 10.64898/2026.09.15.751501

**Authors:** Nicolò Tartini, Lakshmi Niranjana, Helene Gurtner, Ludovico Formenti, Madhav Thakur

## Abstract

Extreme climate events are becoming more frequent, yet their consequences for the recovery of plant–invertebrate relationships under warming remain poorly understood. We investigated over two years of how ground-dwelling invertebrate communities recovered from a 40-days drought in outdoor grassland mesocosms exposed to ambient conditions, constant warming or periodic heatwaves, and both combined. We then examined how post-drought community shifts were associated with above-ground plant biomass, measured concurrently with invertebrate sampling. Drought had no detectable effect on total invertebrate abundance at either recovery time (one or four months after drought). By contrast, diversity showed warming-dependent responses one month after drought and a delayed drought legacy four months later, when diversity declined only under constant warming, with or without periodic heatwaves. At the four-month post-drought sampling, plant biomass–invertebrate diversity coupling was evident only in non-drought plots, with negative relationships under ambient and heatwave regimes but a positive relationship under constant warming alone. This pattern was largely associated with responses of herbivores, particularly Stylommatophora. Our results highlight lagged drought effects in invertebrate diversity that emerged months later and, under warming, extended to weakened plant biomass–invertebrate diversity coupling, pointing to disrupted bottom-up control with potential consequences for food-web functioning.

## Introduction

Invertebrates are the most diverse and abundant group of animals in terrestrial ecosystems. However, their abundance and diversity are declining globally as a consequence of global change drivers, including climate extremes [1–3]. The increasing frequency and intensity of extreme climatic events, including heat waves and prolonged droughts [4,5], pose severe stresses on terrestrial invertebrate populations [6]. For example, drought and heat waves can directly affect invertebrates by altering physiology, life-history traits, and behaviour, while also indirectly affecting resource availability and biotic interactions [1,7–9]. The cumulative effects of these stressors may become severe enough to trigger population breakdowns [3,10], thereby causing strong constraints on their potential recovery [10,11]. Yet, little is known about how invertebrate communities may recover following extreme drought, particularly when in interaction with warming.

The effects of drought on invertebrates lie mostly on desiccation risks and resource availability. For instance, several terrestrial invertebrate groups like orthoptera and bumble bees have been shown to decrease in population and performance during droughts mainly due to, nutrients availability, and starvation [12–16]. Desiccation risk can easily strike Stylommatophora (gastropods) [17], and drought-driven declines following have also been reported in Lepidoptera and sap feeders [7].However, responses are not always uniform across taxa. For instance, groups like Hymenoptera, Diptera, and Hemiptera have been shown to increase in their abundances during and after drought due to increased plant susceptibility and nutritional pulses [18,19]. Indeed, invertebrate responses to drought are likely to depend on other factors, such as on the intensity of stressors [20], reductions in competition, or predation [3]. Invertebrates are particularly responsive to warming because temperature strongly regulates their metabolic rates [8,9,21]. Warming-induced metabolic changes can, in turn, alter development, feeding activity, body size, and voltinism [21–23]. In addition, the extreme temperatures during heat waves usually induce behavioural changes, for example forcing invertebrates to seek shelter in microhabitats [9]. Individuals unable to escape extreme temperatures can suffer physical damage or direct mortality [24,25]. How these acute individual-level effects translate into longer-term community recovery, particularly when warming interacts with drought, remains less clear. Few experiments have tested how drought and warming jointly shape post-drought invertebrate recovery, particularly through repeated assessments capable of capturing transient, delayed, and taxon-specific responses [18,26,27]. Recovery depends on resource availability and favourable conditions, but recurrent droughts or heat waves can accumulate physiological damage, prolonged population impacts, and weaken community recovery [3,10,26,28].

Plants are a major bottom-up resource for most terrestrial invertebrates, but their importance extends beyond food provisioning [29,30]. For instance, many ground-dwelling invertebrate communities depend on vegetation cover or biomass for shelter and microclimate, including using vegetation structure to capture their prey [31–33]. Moreover, greater plant biomass is normally supporting invertebrate abundance across trophic levels, directly influencing herbivores [34,35], and indirectly benefiting predatory and generalist groups [32,36,37]. Furthermore, plants themselves respond to warming and drought, such as changes in productivity, richness, phenology and tissue quality, mediating the climate change effects on invertebrate communities [23,38,39]. For example, constant warming advances phenology, extends the growing season and increases above-ground plant productivity, while decreasing tissue quality [36,40–42]. Post-drought increase in dead plant biomass, can also provide a pulse of resources for several detritivore invertebrates and potentially promoting abundances increases during the recovery phase [18,43]. These changes may strengthen bottom-up effects through greater resource availability. Although, declines in nitrogen content and shift in tissue composition driven by climate warming, can reduce the nutritional quality of plant resources for herbivorous invertebrates, leading to increases in feeding rate and foraging time [9,38,44,45]. These variable responses among different invertebrate groups to warming or drought can cascade to shifts in invertebrate communities, potentially linking post-drought plant biomass recovery to invertebrate recovery through restored bottom-up regulation under warming [31,46].

In this study, we examine how ground-dwelling invertebrate communities recover from extreme drought under different warming regimes, asking whether the recovery patterns are explained by bottom-up control from aboveground plant biomass. For this purpose, we used abundances and diversity metrics to explore the impacts of climate extremes on ground-dwelling grassland invertebrates. In each of the two consecutive years, we crossed an extreme summer drought with constant warming or heatwaves or the combination of both. Furthermore, to account for the temporal dynamics effects in the post-drought recovery, we observed the trajectories at one and four months after drought within a single growing season (figure 1 and figure S1). Within this experimental framework, we tested four hypotheses: (i) Post-drought recovery would vary among taxonomic orders; of the most abundant groups, we expected negative desiccation-driven responses in Stylommatophora, positive resource-driven responses in Hemipterans, and weak responses in generalist multi-trophic groups such as Hymenoptera and Coleoptera. (ii) Warming would delay the recovery of invertebrate diversity following drought, with the strongest and most persistent declines under constant warming combined with heatwaves. (iii) Drought and warming driven shifts on invertebrate diversity would show as changes in invertebrate groups identity turnover, with higher changes under constant warming with heatwaves (iv) Invertebrate diversity or abundance recovery would be partially driven by plant aboveground biomass (bottom-up control; figure 1b).

**Figure 1:**
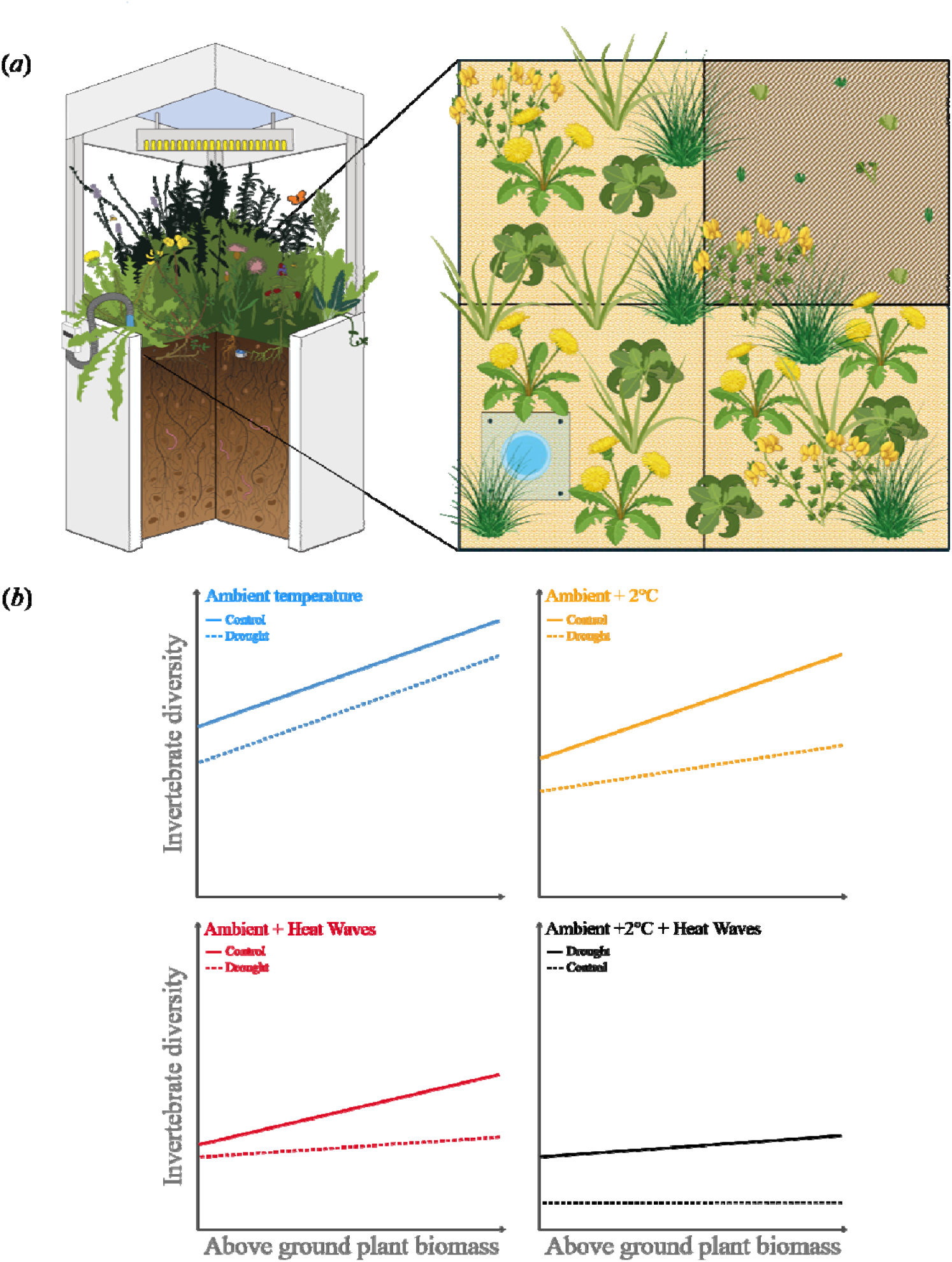
Experimental design and hypotheses. **(a)** Schematic of a mesocosm showing pitfall trap placement. Plants were harvested in one quadrant of each mesocosm, and the pitfall trap was placed randomly in one of the remaining quadrants. The same design was applied in both years of collection (2023 and 2024). **(b)** Expected responses of invertebrate diversity and abundance to aboveground plant biomass across drought treatments and warming regimes. Solid lines denote responses under ambient watering; dashed lines denote responses under drought.

## 2. Materials and Methods

### (a) Experimental set-up

In the years 2023 and 2024, we collected ground dwelling invertebrates using pitfall traps in an outdoor experiment where we simultaneously combined extreme drought and different warming regimes. The experiment is located at the Hasli Ethological Station of the University of Bern, Switzerland (46°58′01.362″ N, 7°23′50.243″ E; 490.6 m a.s.l.). The experiment is composed of 40 semi-natural grassland mesocosms, each covering 1 m², where invertebrate communities established through natural colonization from the surrounding areas (figure 1 and figure S1). The surrounding areas consist of managed meadows, followed by a forest in the south direction, and the river Aare located on the north and west side. The mesocosms are constructed from fibre-reinforced plastic and fitted with metal frames (1 m × 1 m × 0.75 m) supporting infrared heaters and removable roof structures (figure 1a). The drought treatment consisted of an annually repeated extreme drought event, during which all mesocosms were covered with transparent roofs to exclude natural rainfall while avoiding treatment-related differences in light exposure. Half of the mesocosms were assigned to the extreme drought treatment and received no water for 40 days, from the end of May through June, whereas control mesocosms were manually watered according to the regional 30-year precipitation normal [47]. Warming treatments were applied annually from March to October, corresponding to the beginning and end of the growing season in Bern, Switzerland. The warming treatment consisted of four regimes: ambient temperature (Ambient), constant warming at +2°C relative to ambient temperature (Ambient +2°C), periodic heat waves applied monthly for one week at +7°C relative to ambient temperature (Ambient + periodic HW), and constant warming combined with periodic heat waves, reaching +9°C relative to ambient temperature during heat-wave periods (Ambient +2°C + periodic HW; Figure S1a). Warming and drought treatments were randomized and arranged in a fully factorial design, of eight treatment combinations per five replicates. This allowed the assessment of single and combined effects of warming and drought (figure S1a).

The detailed plant communities, substrate, and plant data collection information are provided in Tartini *et al.* (2026). Briefly, the soil substrate consists of a mixture of organic soil, gravel, and sand. To inoculate the mesocosms with microbial and mesofaunal communities, we added a top layer of natural soil collected from the neighbouring grassland. The plant community includes eight native European grassland species (*Holcus lanatus*, *Bromus erectus*, *Trifolium pratense*, *Lotus corniculatus*, *Taraxacum officinale*, *Centaurea jacea*, *Salvia pratense*, and *Prunella vulgaris*) and the invasive North American species *Solidago canadensis*. The plant shoot biomass was collected three times per year in one randomly assigned quadrant (0.25m^2^). Shoot biomass was cut 5 cm above the soil surface, sorted by species, oven-dried at 60°C until full desiccation and weighed. The invertebrate data collection, coincided with the plant collection (figure S1b).

### (b) Invertebrate sampling

Across the two experimental years, we sampled ground-dwelling invertebrate communities using pitfall traps (plastic cylinder, 10 cm height × 5 cm diameter). Invertebrates were collected three times per year: before drought start in late spring (May), one month after drought at the peak of the growing season (late July), and four months after drought in the late growing season (mid-October; Fig. S2). These sampling points were selected to capture shorter- and longer-term post-drought responses within the growing season. Pitfall traps were placed in a randomly selected quadrat within each mesocosm (Fig. 1a). We used one pitfall trap per mesocosm at each sampling point, resulting in 120 trap deployments per year. The vegetation removal quadrant along with the pitfall installation quadrants were also randomly selected, although ensuring that respective sampling occur in different quadrants at every sampling event. The same quadrants for vegetation and pitfall traps were avoided to minimise disturbance from vegetation removal on invertebrates. Moreover, pitfall traps were installed one week before plant clipping. Pitfall traps were filled with 20 ml of a 2:1 propylene-glycol:water solution, with sodium dodecyl sulphate soap (SDS; ReagentPlus®) added to reduce surface tension and improve invertebrate capture efficiency. Rain covers were placed above each pitfall trap to prevent precipitation from diluting or overflowing the preservative solution (Fig. 1a). After seven days, invertebrates collected from each pitfall trap were transferred to 70% ethanol for preservation. In the laboratory, invertebrates were then sorted and identified at the order level using a stereomicroscope (Leica S9i, Leica Microsystems GmbH, Wetzlar, Germany). Once sorted, individuals were counted within each taxonomic group.

### (c) Data analysis

All statistical analyses were carried out in R Statistical Software v.4.3.3 (R Core Team, 2023). For each mesocosm and sampling point, we calculated order richness and Hill numbers using the *renyi* function in the vegan package v.2.7.2 [48]. Specifically, we calculated richness as q = 0, the exponential of Shannon diversity as q = 1, and the inverse Simpson diversity as q = 2. This allowed us to obtain a balanced weighting of the rare and dominant groups [49]. We also analysed total invertebrate abundance and the abundance of the dominant taxonomic groups. To assess changes in community assembly, we calculated rank-abundance curve differences using the package codyn v.2.0.5 [50,51], following an approach similar to Avolio et al. (2019). However, rather than calculating temporal changes in rank-abundance curves, we calculated differences between drought and control treatments within each year. This allowed us to isolate drought-treatment effects rather than temporal turnover. Finally, we investigated invertebrate frequencies to determine whether specific invertebrate orders disappeared under particular climatic treatment.

Models were successfully fitted for four groups: Hemiptera, Hymenoptera, Stylommatophora and Coleoptera which together accounted for 79.70% of all individuals (figure S3). The models for the remaining groups, representing 20.30% of individuals, failed to converge by data rarefication. Therefore, we aggregated the abundances of the latter groups and fitted a model to the pooled abundance data. We fitted each taxa specific abundance and diversity metric as a response variable in generalized linear mixed-effects models (GLMMs) using glmmTMB v.1.1.9 [52]. Fixed effects included warming regimes, drought treatment and sampling points, all fitted as categorical variables in a full three-way interaction (Warming × Drought × sampling points). Furthermore, the year of collection was included as additive covariate while the plot identity was included as a random effect.

For the pre-drought sampling point, only data from 2024 was included because spring 2023 preceded the first drought application and therefore drought-treatment effects did not exist at the time. Furthermore, to test if drought influenced the bottom-up interaction within the invertebrates and plant productivity, we fitted another set of models including total above-ground plant biomass as a continuous numerical predictor. To do so, we fitted the models in a full factorial four-way interaction with temperature treatment, drought treatment and sampling point (warming × drought × sampling points x plant biomass). Again, we included the year of collection as a fixed effect and the plot identity was included as a random effect. The fitting of each GLMM was assessed using simulation-based residual diagnostics with DHARMa v.0.4.7 [53]. Error distributions, link functions, and correction procedures for each response variable are reported in table S1.

After fitting the models, we estimated post-drought marginal means, contrasts, regression estimates and effect sizes for both recovery time points, one and four months after drought for all temperature regimes, using emmeans v.1.10.1 [54]. For models including above-ground biomass, marginal and conditional R² values were calculated using MuMIn v.1.48.11 [55]. Post-drought recovery responses were visualised as Cohen’s d with 95% confidence intervals using ggplot2 v.4.0.0 [56]. For models including above-ground biomass, we also visualised predicted regression relationships by reconstructing model slopes for each treatment and recovery time point, with 95% confidence intervals plotted together with the raw data.

## 3. Results

We included 6607 invertebrate individuals across 16 groups in the analysis. The most abundant orders were Stylommatophora 29.79% (n = 1968), Hemipterans 23.81% (n = 1573), Hymenopterans 20.18% (n = 1333), Acari 9.55% (n = 631), Diptera 9.55% (n = 631) and Coleoptera 5.93% (n = 392), the remaining 10 groups accounted for 1.20% (n = 79) of total abundance (details in supplementary figures 3 and 4, supplementary table 2).

### (a) Post-drought recovery of invertebrate abundance and diversity

One month after drought, total invertebrate abundance decreased under ambient temperature (*β* = -0.706, *z* = -2.113, *p* = 0.035) but not under any warming regime (figure 2a, table S3). Total abundance recovered to control levels under all treatments four months after drought (table S3). By contrast, invertebrate diversity recovery after drought was more pronounced under warming regimes (figure 2b–d). Four months after drought, order richness remained significantly reduced under constant warming with periodic heat waves (*β* = -0.808, *z* = -2.157, *p* = 0.031; figure 2b). Shannon diversity was also significantly reduced at this recovery stage under both constant warming (*β* = -0.878, *z* = -2.191, *p* = 0.028) and constant warming with periodic heat waves (*β* = -0.876, *z* = -2.184, *p* = 0.029). Simpson diversity showed a similar pattern, with significant reductions under constant warming (*β* = -0.920, *z* = -2.289, *p* = 0.022) and constant warming with periodic heat waves (*β* = -0.805, *z* = -2.001, *p* = 0.045).

**Figure 2:**
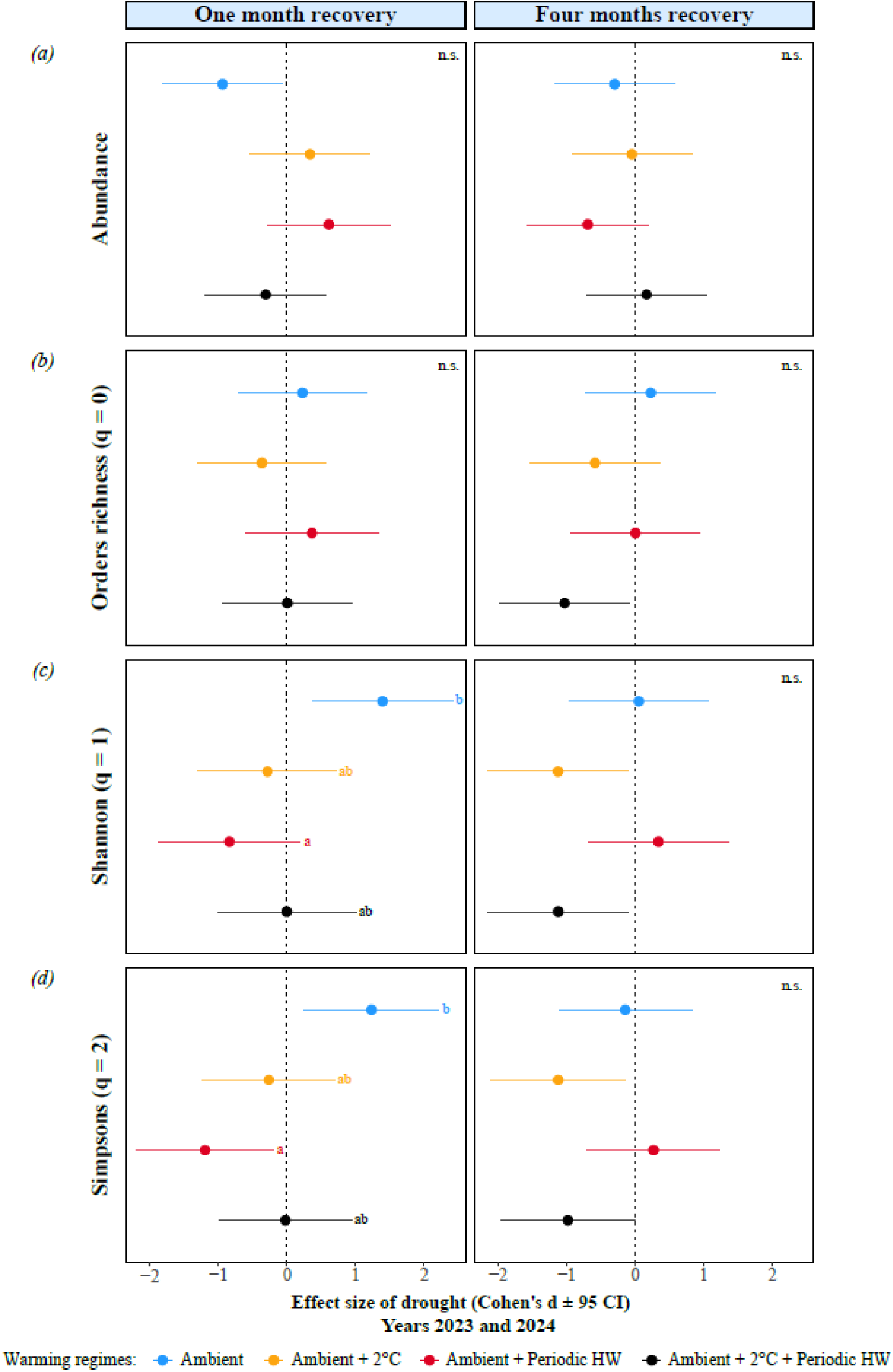
Invertebrate abundance and diversity one and four months after drought under each warming regime across two years of data collection (2023 and 2024). Post-drought recovery is shown for total abundance and for three diversity indices at the order level (Hill numbers: richness, Shannon and Simpson, the latter as a measure of evenness). Values are Cohen’s d with 95% confidence intervals (CIs). Drought effects are significant where CIs do not overlap zero. Significant differences between warming regimes are indicated where present, and labelled non-significant (n.s.) otherwise.

Among individual orders, Stylommatophora and Hymenoptera showed incomplete recovery one month after drought (figure S6). Hymenoptera abundance remained significantly reduced under ambient temperature (*β* = -1.019, *z* = -2.701, *p* = 0.007; table S7), whereas Stylommatophora showed significant post-drought under-recovery under periodic heat waves (*β* = -1.316, *z* = -2.443, *p* = 0.015; table S8). In contrast, Hemiptera and Coleoptera showed no detectable responses to drought (tables S9-S11). Four months after drought, Stylommatophora was the only order that still showed a significant drought response, with abundance remaining reduced under periodic heat waves (*β* = -0.833, *z* = -2.541, *p* = 0.011; figure S6).

### (b) Community assembly after drought

Rank-based turnover in invertebrate groups between drought and control plots was already present one month after drought, ranging from 0.23 to 0.19, with the lowest difference respect to ambient conditions under constant warming combined with periodic heat waves (*β* = -0.039, *z* = -2.651, *p* = 0.023; figure 3a and figure S7). This turnover increased significantly by four months after drought (*β* = 0.033, *z* = 4.413, *p* < 0.001, figure 3a). Rank-abundance curve differences followed a different trajectory instead: one month after drought, curve differences showed only marginal differences between control and drought plots (figure 3b). By four months, curve differences had decreased significantly for invertebrate communities under ambient (*β* = 0.425, *z* = -4.074, *p* < 0.001), periodic heat waves (*β* = 0.365, *z* = -4.668, *p* < 0.001), and constant warming with periodic heat waves (*β* = 0.653, *z* = -2.027, *p* = 0.043), while under constant warming alone no significant temporal difference was observed (*β* = 0.914, *z* = -0.426, *p* = 0.670; figure 3b).

**Figure 3:**
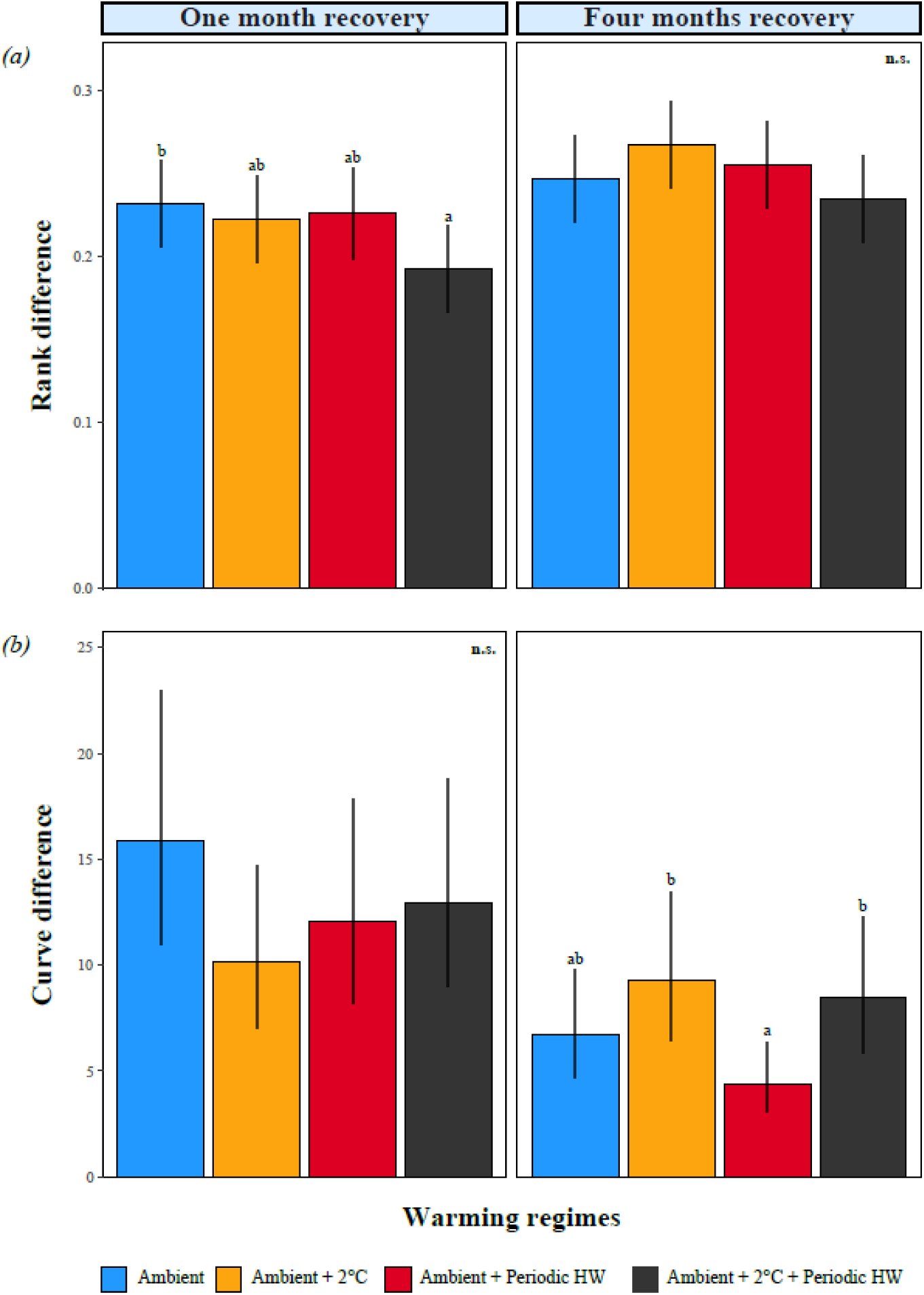
Rank and curve differences between control and drought. We display the change in the order rank and curve depending on the drought treatment and the warming regimes. The bar result from the model output (emmeans) and they represent the differences between drought and control plots. The significance withing warming regimes, are obtained from post-hoc contrast from glmms. The grouping on the warming regimes are displayed when present or labelled as non-significant (N.s.) if no effect was observed.

### (c) Plant above-ground biomass and invertebrate diversity

Plant above-ground biomass did not respond to drought and returned to undisturbed condition under all treatments already one month after the drought (figure S8). At that point of recovery, plant above-ground biomass was not related to invertebrates (figure 4). Among all the diversity metrics and treatments, we observed only one significant relationship: richness increased under ambient control conditions at one-month recovery stage (*slope* = 0.011, *z* = 2.341, *p* = 0.019; figure S9; tables S14–S18). By four months after the drought, biomass became a significant predictor for every diversity metric (figure 4b, figure S10, table S14). For Simpson diversity, the shift was clearest: it increased with biomass under constant warming (*slope* = 0.005, *z* = 2.710, p = 0.007), but declined with biomass under ambient conditions (slope = -0.002, z = -1.976, p = 0.048), periodic heat waves (*slope* = -0.003, *z* = -2.038, *p* = 0.042) and constant warming with periodic heat waves (*slope* = -0.004, *z* = -3.281, *p* = 0.001; figure 4b). For invertebrate abundance, the pattern was similar: it increased with above-ground biomass under every treatment except constant warming, where no relationship with biomass was detected (figure S9). For richness and Shannon, biomass was likewise a positive predictor under constant warming (*slope* = 0.018, *z* = 2.392, *p* = 0.017; and *slope* = 0.016, *z* = 3.306, *p* < 0.001, respectively), while exponential Shannon decreased with biomass under constant warming combined with periodic heat waves (*slope* = -0.009, *z* = -2.425, *p* = 0.015; figure S10). Group-specific abundance–biomass relationships are reported in detail in the appendix (figures S11–S12; tables S19–S22).

**Figure 4:**
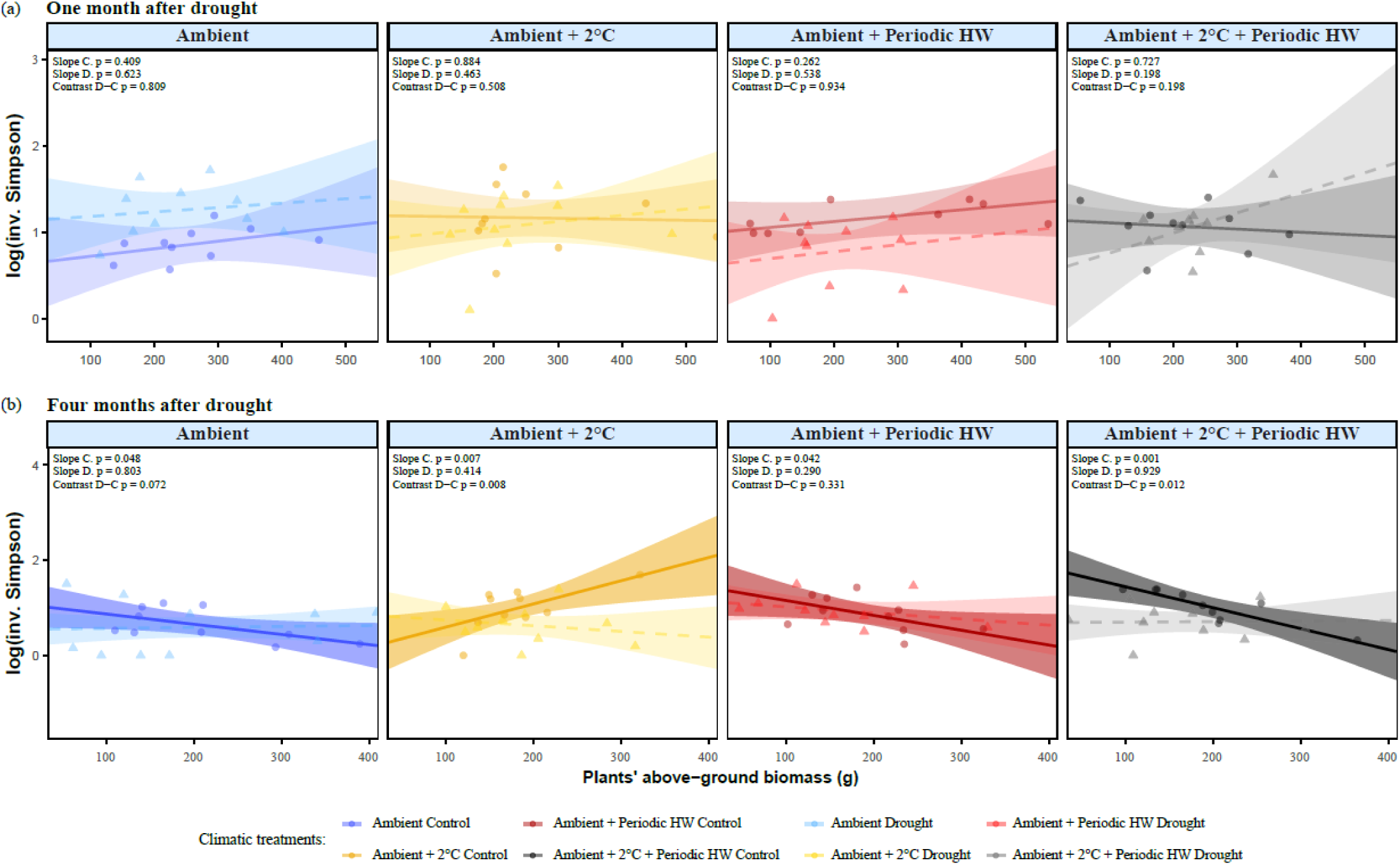
Relationship between Simpson diversity (Hill number q = 2) and aboveground plant biomass. (a) One month after drought and (b) four months after drought. Lines are linear regressions fitted from generalised linear mixed model outputs, with shaded areas showing 95% confidence intervals (CIs). Solid lines denote significant slopes; dashed lines denote non-significant slopes. Circles are ambient-watering plots, triangles are drought plots. Slope significances and the significance of contrasts between slopes are shown for all regressions. C, ambient watering; D, drought; HW, heatwave. Equivalent figures for the remaining diversity indices are given in Supplementary Figures S9 and S10.

## 4. Discussion

Over two years, we tested how ground-dwelling invertebrate abundances recover from extreme drought and how warming shapes this recovery. Even if invertebrate abundances recovered rapidly overall, drought disrupted the link between invertebrates and above-ground plant biomass, leading to a breakdown of bottom-up regulation. Drought also impacted invertebrate diversity, with lasting effects on population composition under constant warming, with and without heatwaves, detectable even four months after the drought. These results match our hypotheses: extreme drought not only impacts invertebrate abundance or diversity, but also weakens the link between invertebrates and plants. The relationship with plant biomass, however, did not match our predictions: it was negative, and emerged only during the later phase of recovery (figure 1), suggesting that bottom-up regulations of invertebrate communities under recovery after extremes are likely to be disrupted.

We hypothesized that invertebrate abundances and diversity and above-ground biomass would have a positive relationship, as well as that drought would weaken the link between invertebrates and biomass. While drought effectively led to a bottom-up regulation breakdown, this only emerged in the later recovery, where plant biomass-invertebrate links were present only in water control mesocosms. Of which, the positive link of invertebrate abundances and plant biomass (Figure S10), aligned to our expectations and with previous studies, which reported the same pattern between plant productivity and invertebrate abundance [32,37,57]. However, the negative invertebrate diversity–plant biomass relationship in regularly watered systems suggests that higher plant productivity favoured a few invertebrate orders, which increased their overall dominance. These results only partially supports our hypotheses: abundance increased with biomass, but diversity decreased. Moreover, such response patterns cannot be attributed to a drought-driven decline in plant productivity. Indeed, above-ground biomass recovered rapidly after drought at both total- and species-level biomass, as reported for the same experiment [58]. The negative diversity–biomass relationships, can be explained by the response of dominant groups such as Stylommatophora, Hemiptera and Hymenoptera, which showed positive relationships with biomass mainly in water control plots (figures S11–S12). These results confirm that drought weakened the link between these dominant groups and plant biomass.

While drought explained the breakdown in bottom-up regulation, invertebrate abundance, meanwhile, followed a more complex pattern: firstly, its recovery was not always complete, with under-recovery at ambient conditions. Secondly, contrary to our hypotheses, the negative abundance response to drought was not further affected by warming (figure 2 and figure 4). One likely explanation is that warming independently suppressed certain invertebrate populations, narrowing the contrast between drought treatments. This was indeed the case for Hymenoptera and Stylommatophora, which showed warming-induced declines in abundance. These results are aligned with a previous study on invertebrate macro- and meso-fauna where the effect of drought and warming did not interact [18]. Although the negative responses of Hymenoptera contrasted with our expectations, this finding is consistent with the negative effects reported in previous studies [14,59]. Other studies, however, reported positive [18] or neutral effects [60]; such heterogeneity likely reflects substantial variation among species within this diverse insect order. On the other hand, Stylommatophora matched the expected sensitivity of terrestrial gastropods to desiccation [17,61,62].

The under-recovery of Hymenoptera and Stylommatophora also explains why abundance recovery did not correspond to a community composition recovery (figure 2 and figure 3). In fact, the diversity over-recovery observed one month after the drought (figure 2c-d) is not indicative of a positive response: it was driven by declines in the dominant groups (figure S6). Declines of abundant taxa are increasingly recognized as a common observation in insects [3], and recent large-scale syntheses show that these declines can themselves produce artefactual increases in diversity indices [11]. A similar pattern has been proposed for Hymenoptera specifically, where shifts in community-level metrics can be generated by small changes in the abundance of the ecologically dominant Formicidae (ant) family [60]. By contrast, drought reduced diversity over the same period under periodic heat waves, suggesting only a subset of taxa maintained or increased their dominance (figure 2b-d), consistent with previous findings in Amazonian ants where drought decreases overall richness while generalist and thermophilic species specifically increased in frequency [59]. The presence of such shifts in taxon abundances and diversity metrics (Figures 2 and 3), support our hypothesis that drought leads to rank shifts in order abundances.

The appearance of delayed effects in the later phase of recovery (figure 2 and figure 4) shows how the impacts of extreme events can be overlooked with single point assessments. For instance, the effects of warming can change according to the season, with no effects in summer while being present in the cooler ones [63]. Although, the lagged responses did not only reveal a seasonal effect. They likely reflect several mechanisms acting on different invertebrate groups, which differ in their sensitivity to drought, warming and post-drought resource availability [13,64]. Warming, for example, may have directly favoured thermophilic taxa, while, heatwaves and reduced temperature buffering due to canopy dieback can impose acute stress on heat-sensitive invertebrates [9,14,65,66]. This responses partially support our hypothesis that warming would slow post-drought recovery. Moreover, during drought, herbivores faced reduced food availability [61]. During the recovery phase, above-ground plant biomass rapidly regrew, but leaf traits did not [58], a combination that may have made aerial plant tissues more palatable and so benefited some herbivorous groups during recovery [67,68]. Altogether, these pathways likely pushed taxa in opposite directions across feeding guilds, leading to the observed short-term and delayed effects on community diversity, with drought effects surfacing at different time points (figures 2 and 3).

## Conclusions

Our results show that drought and warming reshape ground-dwelling invertebrate communities during post-drought recovery, disrupting bottom-up regulation by plant biomass and richness, eroding the dominance of key taxa, and generating responses that emerge only later in the growing season. These dynamics are of growing importance in a world where anthropogenic climate change will intensify the frequency and strength of extreme climate events [4]. The consequence for grasslands is that numerical recovery outpaces functional recovery: abundance and richness may return towards pre-drought levels while the interactions linking plants to consumers remain uncoupled, leaving the community reassembled rather than restored. Detecting this required repeated sampling. A single post-drought survey would have recorded recovery and missed the delayed and compositional shifts underlying it. Given the scarcity of experimental studies which follow invertebrate communities beyond the immediate aftermath of an extreme event [1,39,69], we encourage long-term designs that track recovery across seasons and years. Future work should test whether the plant–invertebrate decoupling we observe extends to canopy-associated invertebrate taxa, and whether resolving feeding guilds clarifies how drought alters the transfer of plant biomass and resource quality to herbivores.

## Acknowledgments

We thank, Felix Rentschler, Anine Wyser, for assisting with the data collection and invertebrate sorting.

## Author contribution

MPT conceived the study. NT, HG, LN and LF collected, sorted and counted the invertebrates. NT analysed the data with substantial input from MPT. NT and MPT wrote the first draft of the manuscript. All authors contributed to revising the manuscript.

## Declaration of AI use

We used AI-assisted tools (Claude, Anthropic) to correct grammar, to improve language clarity, and to assist for support in R data visualization. All other aspect of the study are the authors’ own.

