## Supplementary Figures and Tables for "Extreme drought disrupts the bottom-up regulation of ground-dwelling invertebrate diversity under climate warming"

**Supplementary information**

**Extreme drought disrupt Plant biomass relationship with invertebrate diversity across warming regimes**

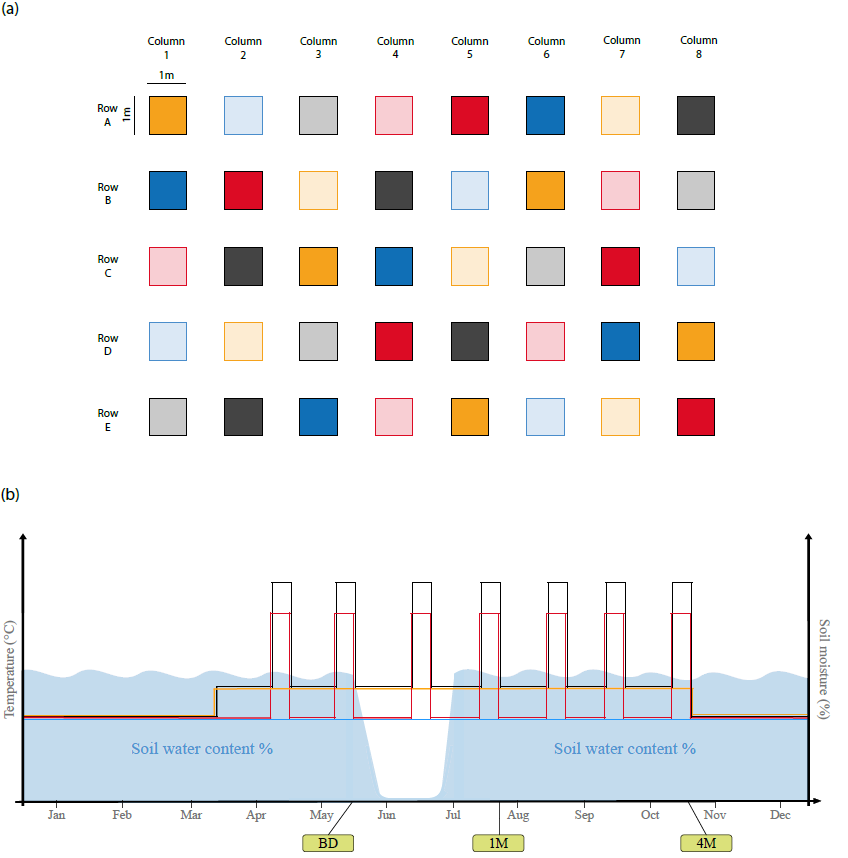
Figure S1: Mesocosm set up and experimental design. The design if fully factorial. (a) Arrangement of the mesocosm and treatments. In blue ambient temperature, in orange constant warming (+2°C), in red periodic heatwaves (+10°C), in black combination of constant warming (+2°C) and periodic heatwaves (+10°C). Lighter colour represent plots where drought is being applied. (b) Seasonal dynamics of the treatments. Constant warming starts in early march and it stops at the end of October. The heatwaves are applied one time per month for one week for a total of 7 heatwaves. Drought is applied in the end of may for almost all the month of June. BD = Data collection before the drought, 1M = data collection one month after the drought, 4M = data collection four month after the drought

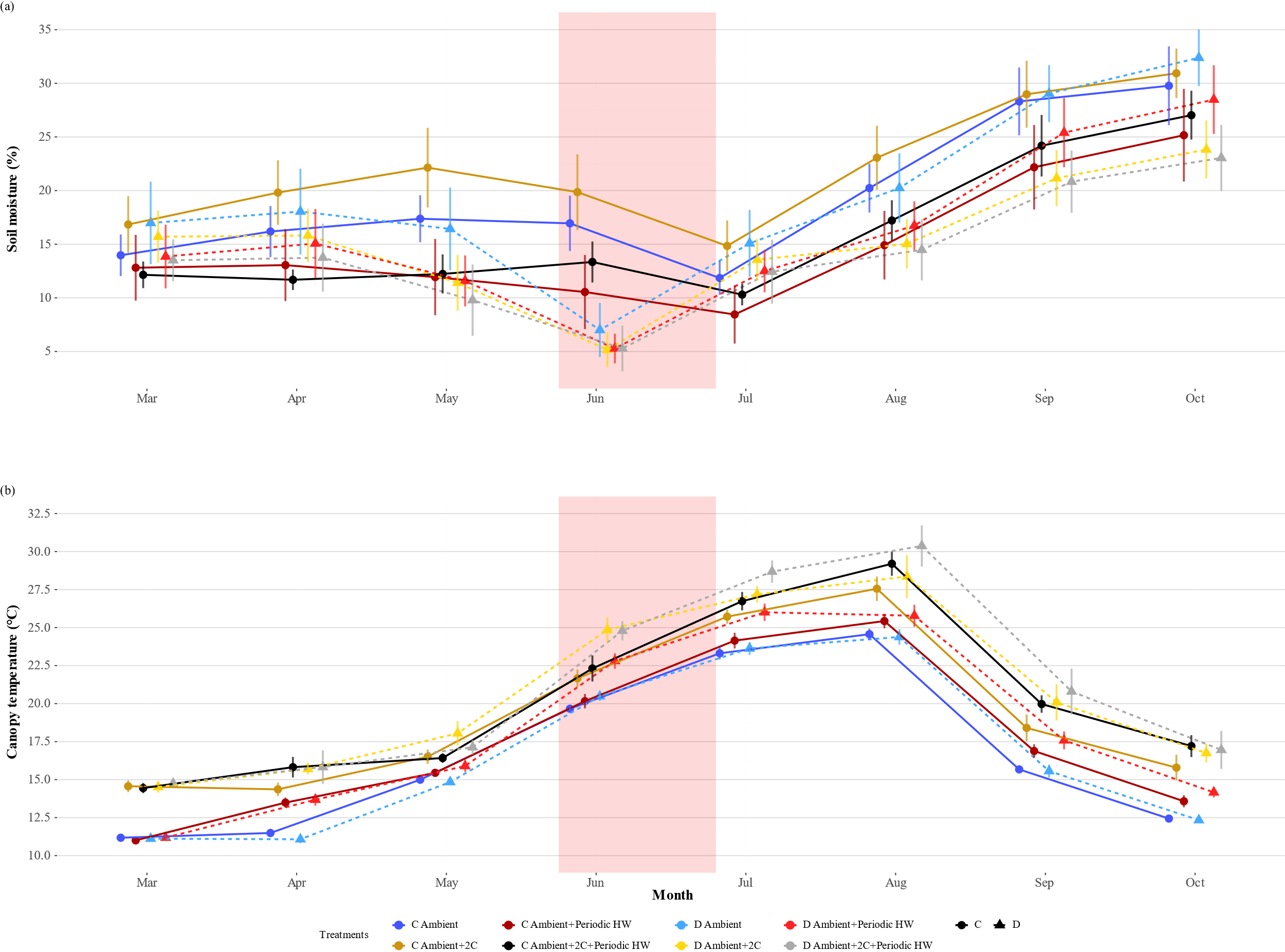

Figure S2: Monthly averages of soil Moisture and canopy temperature measurement taken in the mesocosms. For both moisture and temperature we display the mean ± SE of the growing season (March-Orctober). (a) Soil moisture monthly average according warming and drought treatments,. The Soil moisture was measured via multidepth sensors in each mesocosm. (b) Monthly average of the Air temperature in the canopy. Measurements were taken in the centre of each mesocosm 15 cm above the soil level. The colours correspond to the warming regimes. Circles and Solid lines represent water control plots, triangles and dashed lines represend dorught plots. The read area correspond to the period where drought was applied. C = Control, D = Drought.

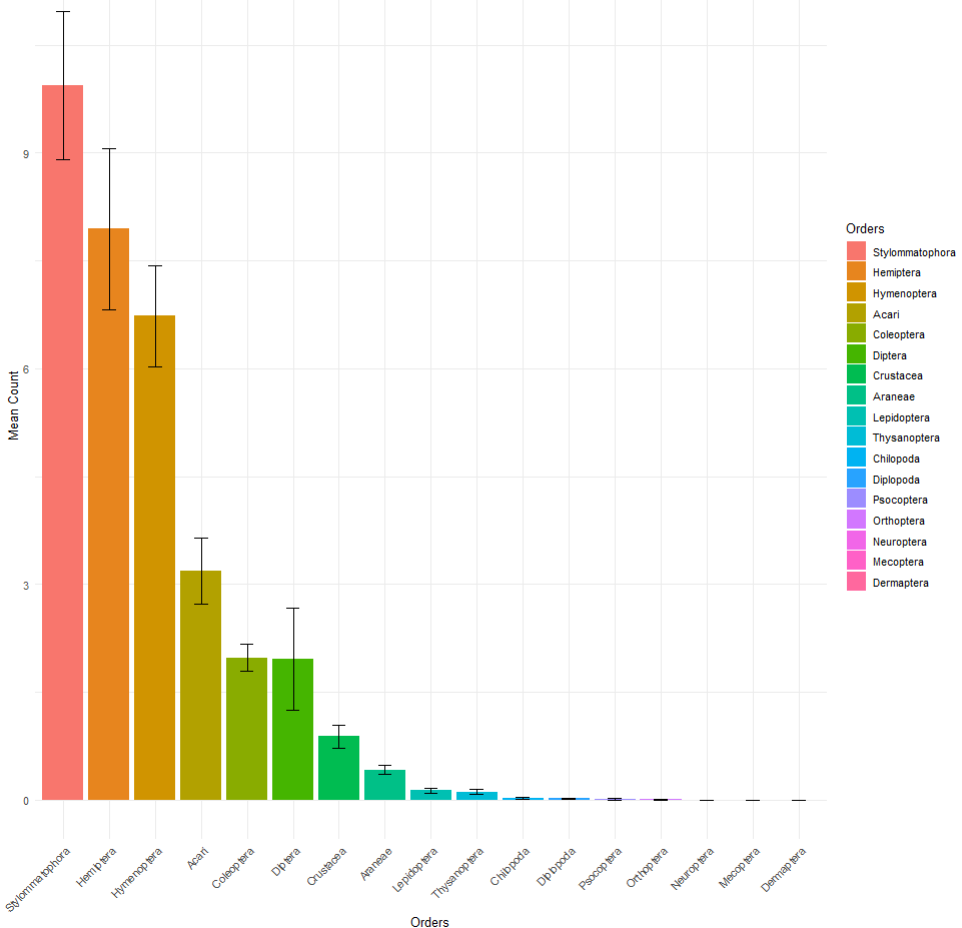

Figure S3: Total orders abundance of the whole data collection(yeas 2023-24). The data represent the mean counts per order found in the mesocosm. The error bars are the standard error (SE). The groups are sorted form the most abundant to the less abundant ones.

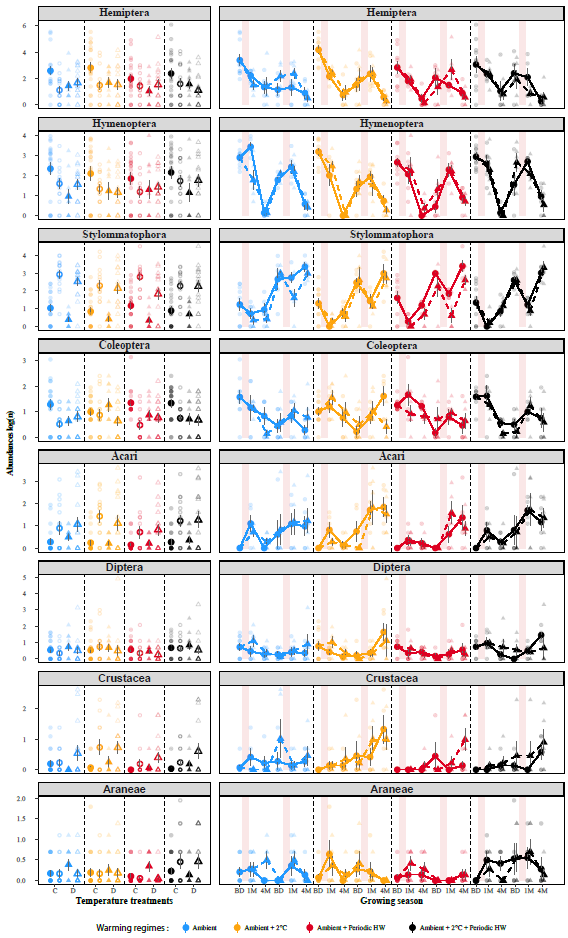

Figure S4: Abundances and average means of the major observed orders. The left panels represent the average abundances of the main Orders in 2023 and 2024 at different warming regimes according the water treatments. The right panels represent the abundances dynamics across the sampling time. The values in both left and right panels are log scaled. The light red surface in the second panels correspond to the drought period. Circles points are referred to water control, triangles to drought plots. Full circles and Triangles = year 2023, Hollow circles and tringles = year 2024. BD = Before the drough, 1M = One month after the drought, 4M = Four month after the drought.

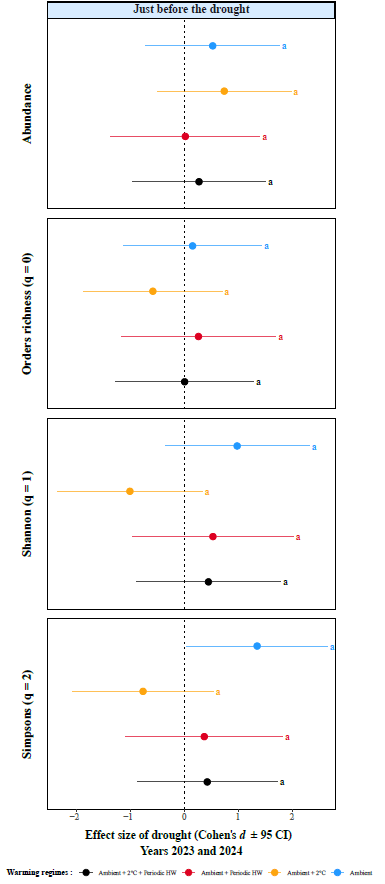

Figure S5: Invertebrates’ abundance and diversity before the drought per each warming regime. We show the legacy effect of drought in for the total population abundances, and three diversity indices at the order scale (total abundance, orders richness q = 0, Shannon q = 1, and Simpson q = 2). The values are Cohen’s d 95% confidence intervals (CIs). Drought effects are significant when the Cis do not overlap with zero. Significance differences within warming regimes are displayed with letter, no statistical effect within the warming regimes was observed.

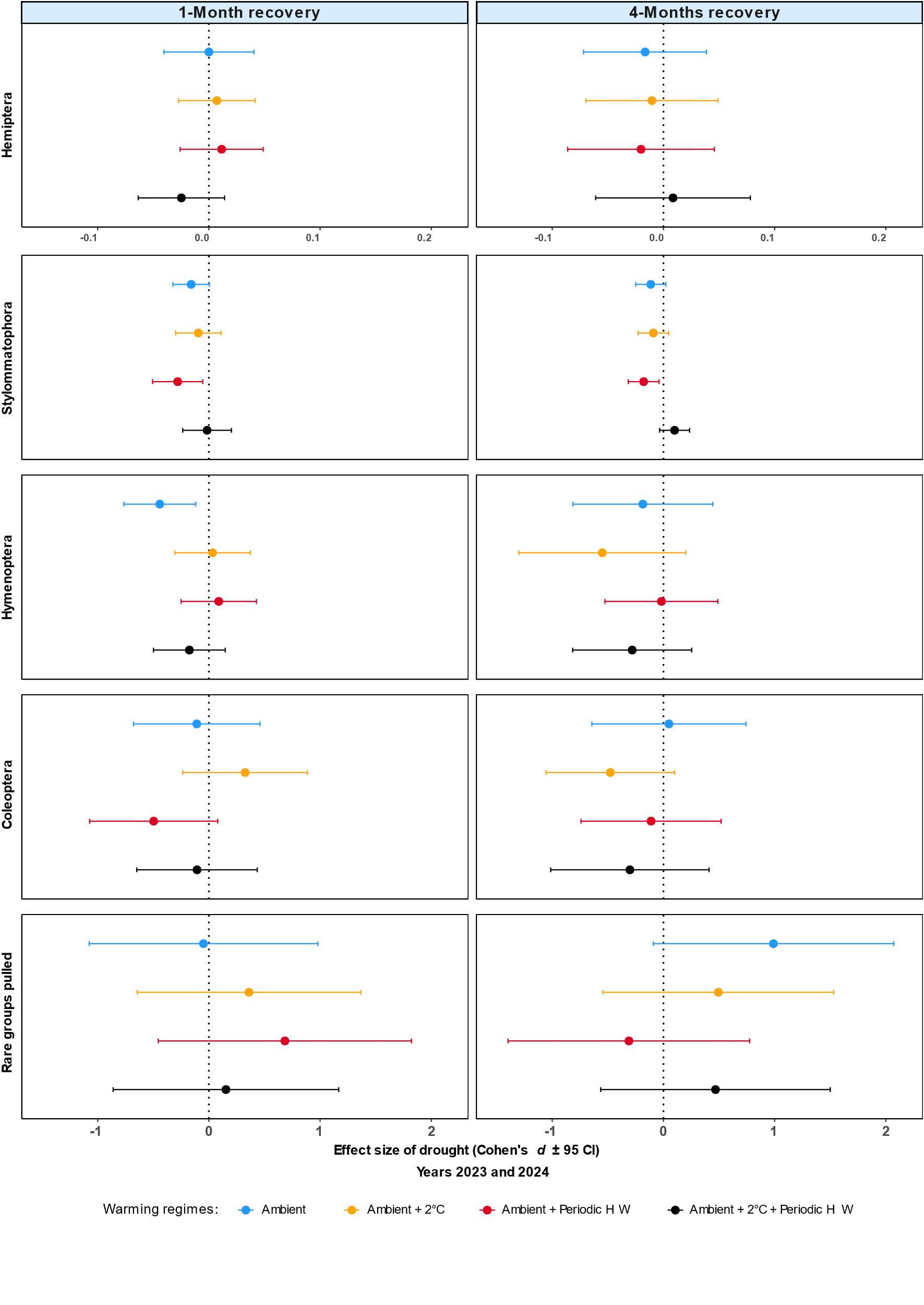

Figure S6: Effect of Drought on the main invertebrates groups at one and four months and four months of recovery. We show pot-drought recovery across four different warming regimes for four more representative orders. The values are Cohen’s d 95% confidence intervals (CIs). Drought effects are significant when the Cis do not overlap with zero.

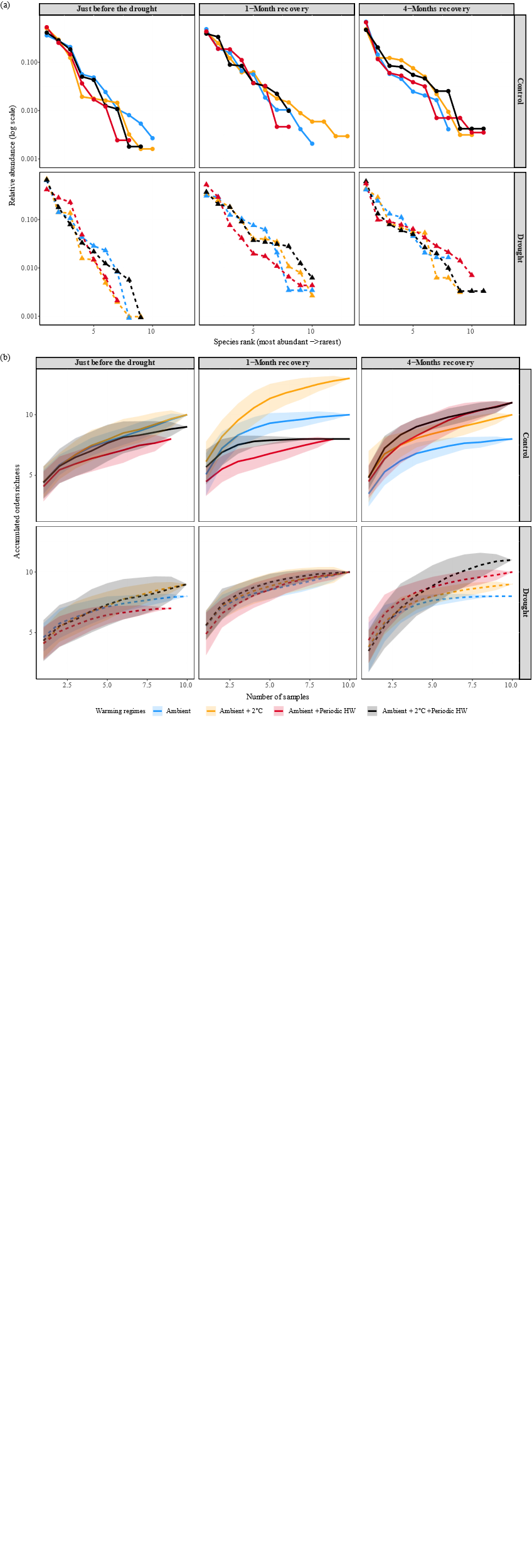

Figure S7: Rank abundances curves. (a) Relative abundance curve according the warming and water regimes. For each sampling point, the abundances are ranked form the most abundant to the less abundant orders. Values are log scaled in reason to ease the visualization. (b) Cumulative order richness according the warming and water regimes. For each sampling point, we display the cumulative order richness the permutation shown are obtained from plot having the same sampling point, warming and water regimes.

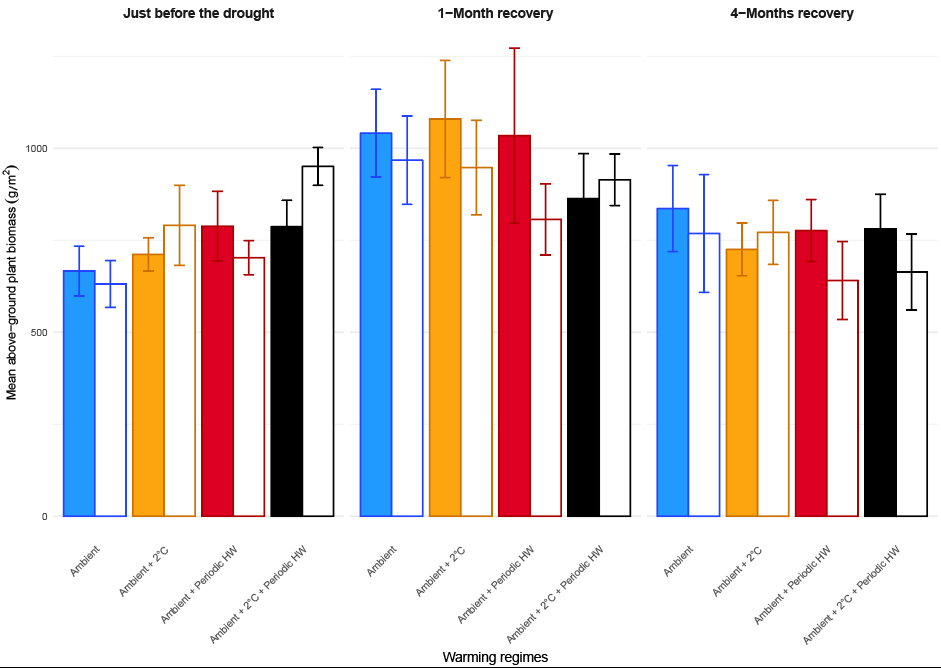

Figure S8: Above ground plant biomass. (b) Average above-ground plant biomass per treatment and sampling point. Biomass is expressed ad gramps per square meter. Bar with colour filling are water control condition, hollowed bars correspond to plots submitted to early-summer drought. The error bar are expressed via standard error.

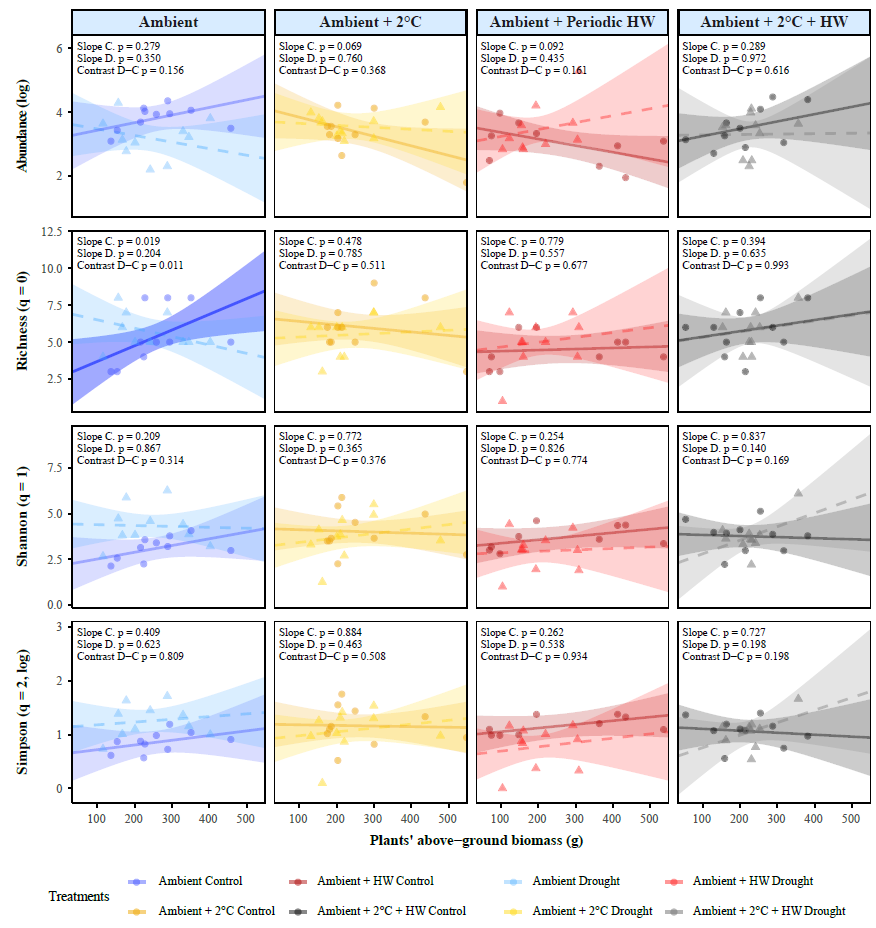

Figure S9: Regression of the studied diversity indexes, Total counts, richness, Shannon (q=1) and Simpson q=2) and plant biomass at 1-Month recovery data point. We show the linear regressions between invertebrate diversity indexes and the above-ground plant biomass, obtained from the outputs of the generalized mixed effect models. The areas of the regressions represent the 95% confidence intervals (CI). Solid lines represent significant regressions dashed lines represent the non-significant ones. Circles points represent water control values, Triangles represent the drought ones. We display the significances of the slopes of the regressions as well the significance of their contrasts for all the slopes. Abbreviations: C = Water control plots, D = water drought plots, HW = Heat Waves.

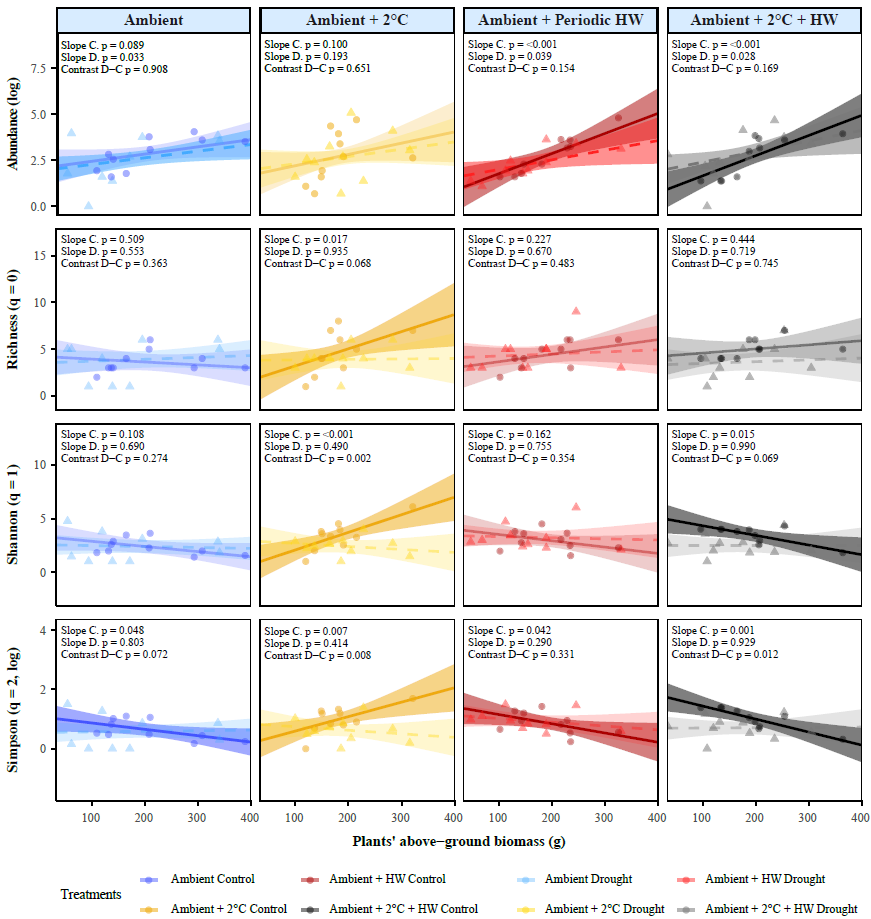

Figure S10: Regression of the studied diversity indexes, Total counts, richness, Shannon (q=1) and Simpson q=2) and plant biomass at 4-Months recovery data point. We show the linear regressions between invertebrate diversity indexes and the above-ground plant biomass, obtained from the outputs of the generalized mixed effect models. The areas of the regressions represent the 95% confidence intervals (CI). Solid lines represent significant regressions dashed lines represent the non-significant ones. Circles points represent water control values, Triangles represent the drought ones. We display the significances of the slopes of the regressions as well the significance of their contrasts for all the slopes. Abbreviations: C = Water control plots, D = water drought plots, HW = Heat Waves.

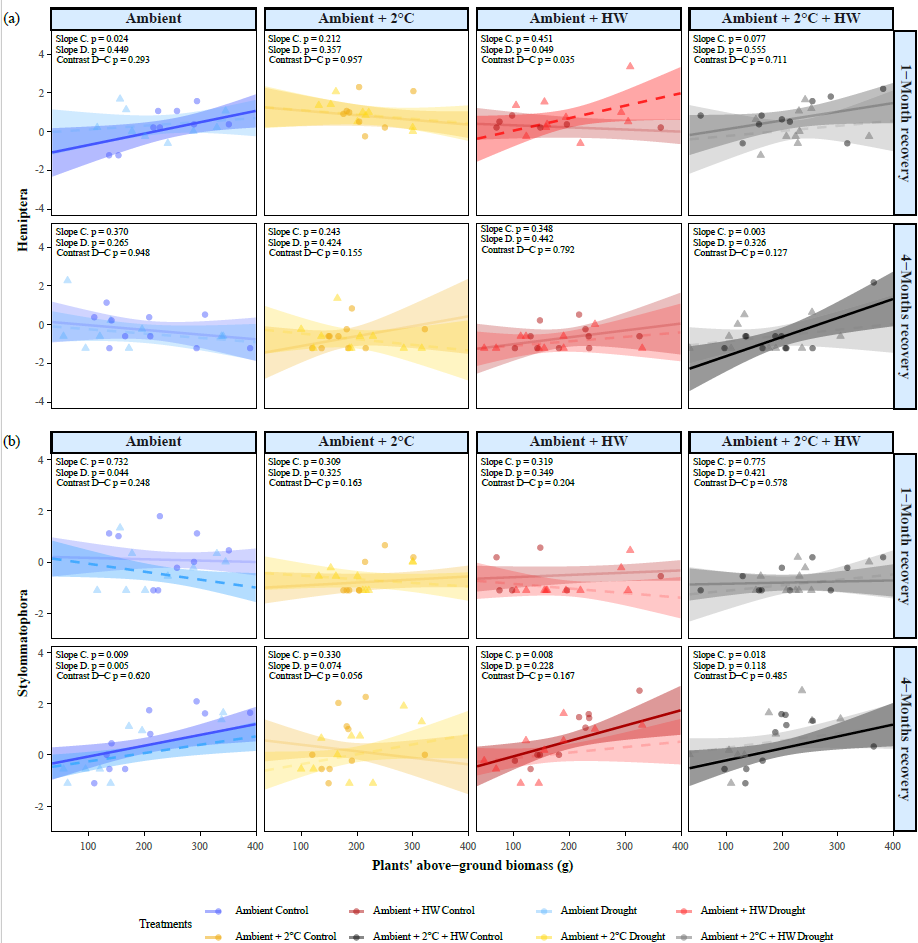

Figure S11: Regression of the sap feeders and herbivore orders (Hemiptera and Stylommatophora). We show the linear regressions between two orders abundances and the above-ground plant biomass. The regressions are obtained from the outputs of the generalized mixed effect models. The areas of the regressions represent the 95% confidence intervals (CI). Solid lines represent water control regressions dashed lines represent the drought ones. Circles points represent water control values, Triangles represent the drought ones. We display the significances of the slopes of the regressions as well the significance of their contrasts for all the slopes. Abbreviations: C = Water control plots, D = water drought plots, HW = Heat Waves.

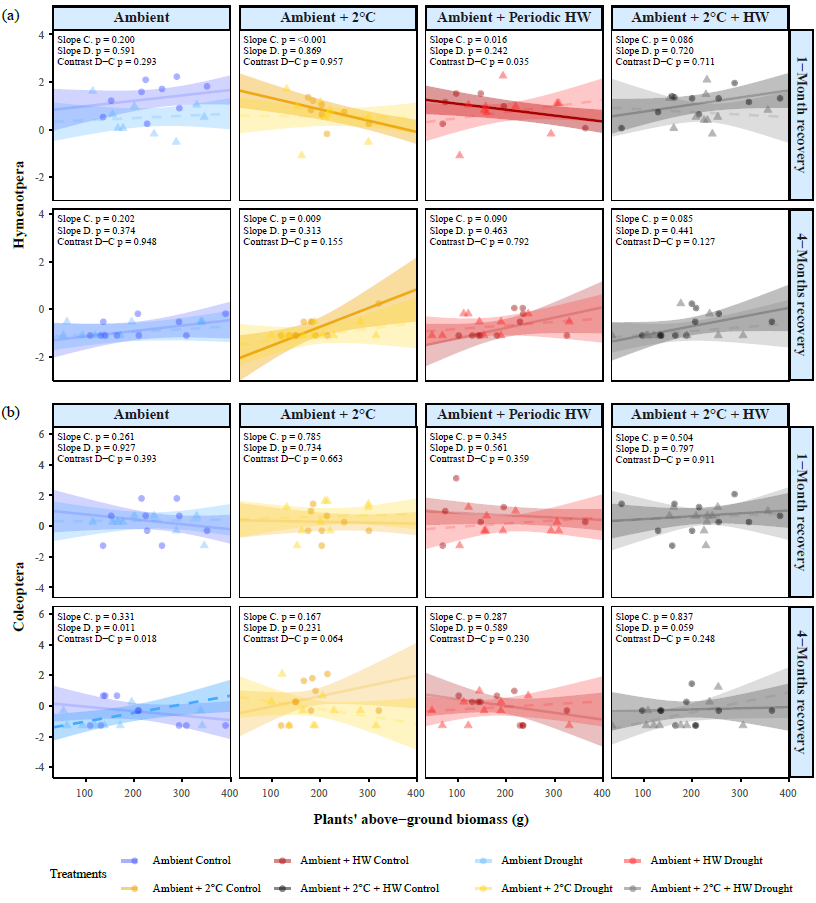
 Figure S12: Regression of Hymenoptera and Coleoptera. We show the linear regressions between two orders abundances and the above-ground plant biomass. The regressions are obtained from the outputs of the generalized mixed effect models. The areas of the regressions represent the 95% confidence intervals (CI). Solid lines represent water control regressions dashed lines represent the drought ones. Circles points represent water control values, Triangles represent the drought ones. We display the significances of the slopes of the regressions as well the significance of their contrasts for all the slopes. Abbreviations: C = Water control plots, D = water drought plots, HW = Heat Waves.

Table S1: Summary of fitted model specifications for the diversity index and abundances analyses . We include models with or without aboveground plant biomass regression as explanatory variable. For each kind of model and response variable we show corrections, scaling and used distribution, and whenever zero inflation was applied. Abbreviations: Zi formula: Zero inflation formula.

| Model | Response variable | Center scaling | Log correction | Distribution | Zi. formula |
| --- | --- | --- | --- | --- | --- |
| Effect of experimental treatments | Order richness (q=0) | Yes | No | Gaussian | No |
|  | Shannon (q=1) | Yes | No | Gaussian | No |
|  | Simpson (q=2) | Yes | Yes | Gaussian | No |
|  | Rank difference | No | No | Gaussian | No |
|  | Curve difference | No | Yes | Gaussian | No |
|  | Total abundances | Yes | Yes | Gaussian | No |
|  | Hemiptera | No | No | Genpois | No |
|  | Stylommatophora | No | No | nbinom2 | ~1 |
|  | Hymenoptera | No | No | nbinom2 | ~1 |
|  | Coleoptera | No | No | nbinom2 | ~1 |
|  | Rare groups | No | No | nbinom2 | No |
| Including biomass regression | Order richness (q=0) | No | No | Gaussian | No |
|  | Shannon (q=1) | No | No | Gaussian | No |
|  | Simpson (q=2) | No | Yes | Gaussian | No |
|  | Total abundances | No | Yes | Gaussian | No |
|  | Hemiptera | Yes | Yes | Gaussian | No |
|  | Stylommatophora | Yes | Yes | Gaussian | No |
|  | Hymenoptera | Yes | Yes | Gaussian | No |
|  | Coleoptera | Yes | Yes | Gaussian | No |

Table S2: Invertebrates abundance summary by order, year, and season. We show the abundance of each observed invertebrate order, by year and timepoint of collection as well as their relative total number of observations. Abbreviations: JBD: Just before the drought, 1M. One-month after the drought recovery, 4M: Four-months after the drought recovery

|  | | **2023** | | **2024** | | |  |
| --- | --- | --- | --- | --- | --- | --- | --- |
| **Order** | **JBD** | **1M** | **4M** | **JBD** | **1M** | **4M** | **Total** |
| Coleoptera | 142 | 151 | 57 | 36 | 78 | 70 | 534 |
| Hemiptera | 2703 | 344 | 171 | 438 | 572 | 48 | 4276 |
| Diptera | 64 | 64 | 29 | 12 | 27 | 256 | 452 |
| Mecoptera | 6 | 0 | 0 | 0 | 0 | 0 | 6 |
| Araneae | 5 | 24 | 11 | 19 | 22 | 7 | 88 |
| Acari | 0 | 49 | 11 | 126 | 257 | 188 | 631 |
| Crustacea | 1 | 7 | 12 | 53 | 31 | 73 | 177 |
| Hymenoptera | 848 | 619 | 6 | 272 | 388 | 48 | 2181 |
| Lepidoptera | 2 | 6 | 11 | 0 | 7 | 2 | 28 |
| Neuroptera | 4 | 0 | 0 | 0 | 0 | 0 | 4 |
| Stylommatophora | 199 | 16 | 67 | 671 | 202 | 1012 | 2167 |
| Thysanoptera | 0 | 1 | 0 | 2 | 16 | 4 | 23 |
| Chilopoda | 0 | 0 | 0 | 1 | 2 | 2 | 5 |
| Orthoptera | 0 | 1 | 0 | 1 | 0 | 0 | 2 |
| Diplopoda | 0 | 0 | 0 | 0 | 2 | 2 | 6 |
| Psocoptera | 0 | 0 | 0 | 0 | 1 | 2 | 5 |
|  |  |  |  |  |  |  | 10581 |

Table S3: Estimated marginal means of drought by temperature and seasons on the invertebrate abundances. Output of the linear mixed-effects models testing invertebrate total abundances to warming, watering and season. The model included water treatment, warming regimes, harvest time point as explanatory variables. The marginal means were calculated intercepting the water treatment, while the significant estimates were obtained contrasting the differences form water and drought plots (Drought minus Control). Significant *P. values* are highlighted in bold. Abbreviations: Emmean = Estimated marginal means, SE = Standard errors, *P* = P. values, Amb = Ambient temperature; +2 °C = Ambient +2 °C warming; +HW = Ambient + heatwave warming (+10 °C); +2 °C+HW = Ambient +2 °C + heatwave warming; 1-month recovery = One month after drought; 4-months recovery = Four-months after drought.

| Season | Drought Treatment | Warming regimes | emmean | SE | *P* |
| --- | --- | --- | --- | --- | --- |
| Just before the drought | Control | Amb | -23.606 | 2.809 | 0.410 |
|  | Drought | Amb | -23.216 | 2.809 |  |
|  | Control | +2°C | -23.957 | 2.809 | 0.245 |
|  | Drought | +2°C | -23.407 | 2.809 |  |
|  | Control | +HW | -23.731 | 2.814 | 0.986 |
|  | Drought | +HW | -23.722 | 2.814 |  |
|  | Control | +2°C+HW | -23.366 | 2.809 | 0.673 |
|  | Drought | +2°C+HW | -23.166 | 2.809 |  |
| 1-Month recovery | **Control** | **Amb** | **-22.800** | **2.740** | **0.035** |
|  | **Drought** | **Amb** | **-23.506** | **2.740** |  |
|  | Control | +2°C | -23.332 | 2.740 | 0.458 |
|  | Drought | +2°C | -23.084 | 2.740 |  |
|  | Control | +HW | -23.649 | 2.734 | 0.186 |
|  | Drought | +HW | -23.195 | 2.740 |  |
|  | Control | +2°C+HW | -23.135 | 2.740 | 0.482 |
|  | Drought | +2°C+HW | -23.370 | 2.740 |  |
| 4-Months recovery | Control | Amb | -23.823 | 2.740 | 0.497 |
|  | Drought | Amb | -24.050 | 2.740 |  |
|  | Control | +2°C | -23.964 | 2.740 | 0.907 |
|  | Drought | +2°C | -24.003 | 2.740 |  |
|  | Control | +HW | -23.892 | 2.740 | 0.119 |
|  | Drought | +HW | -24.413 | 2.740 |  |
|  | Control | +2°C+HW | -23.994 | 2.740 | 0.719 |
|  | Drought | +2°C+HW | -23.874 | 2.740 |  |

Table S4: Estimated marginal means of drought by temperature and seasons on the invertebrate taxonomic richness. Output of the linear mixed-effects models testing invertebrate total abundances to warming, watering and season. The model included water treatment, warming regimes, harvest time point as explanatory variables. The marginal means were calculated intercepting the water treatment, while the significant estimates were obtained contrasting the differences form water and drought plots (Drought minus Control). Significant p values are highlighted in bold. Abbreviations: Emmean = Estimated marginal means, SE = Standard errors, *P* = P. values, Amb = Ambient temperature; +2 °C = Ambient +2 °C warming; +HW = Ambient + heatwave warming (+10 °C); +2 °C+HW = Ambient +2 °C + heatwave warming; 1-month recovery = One month after drought; 4-months recovery = Four-months after drought.

| Season | Drought Treatment | Warming regimes | emmean | SE | *P* |
| --- | --- | --- | --- | --- | --- |
| Just before the drought | Control | Amb | -17.572 | 2.939 | 0.822 |
|  | Drought | Amb | -17.457 | 2.939 |  |
|  | Control | +2°C | -17.572 | 2.939 | 0.368 |
|  | Drought | +2°C | -18.034 | 2.939 |  |
|  | Control | +HW | -18.270 | 2.944 | 0.726 |
|  | Drought | +HW | -18.069 | 2.945 |  |
|  | Control | +2°C+HW | -17.457 | 2.939 | 1.000 |
|  | Drought | +2°C+HW | -17.457 | 2.939 |  |
| 1-Month recovery | Control | Amb | -16.511 | 2.867 | 0.644 |
|  | Drought | Amb | -16.338 | 2.867 |  |
|  | Control | +2°C | -16.165 | 2.867 | 0.441 |
|  | Drought | +2°C | -16.453 | 2.867 |  |
|  | Control | +HW | -17.024 | 2.861 | 0.462 |
|  | Drought | +HW | -16.742 | 2.867 |  |
|  | Control | +2°C+HW | -16.280 | 2.867 | 1.000 |
|  | Drought | +2°C+HW | -16.280 | 2.867 |  |
| 4-Months recovery | Control | Amb | -17.550 | 2.867 | 0.644 |
|  | Drought | Amb | -17.377 | 2.867 |  |
|  | Control | +2°C | -16.915 | 2.867 | 0.218 |
|  | Drought | +2°C | -17.377 | 2.867 |  |
|  | Control | +HW | -17.088 | 2.867 | 1.000 |
|  | Drought | +HW | -17.088 | 2.867 |  |
|  | **Control** | **+2°C+HW** | **-16.742** | **2.867** | **0.031** |
|  | **Drought** | **+2°C+HW** | **-17.550** | **2.867** |  |

Table S5: Estimated marginal means of drought by temperature and seasons on Shannon’s diversity (q=1). Output of the linear mixed-effects models testing invertebrate total abundances to warming, watering and season. The model included water treatment, warming regimes, harvest time point as explanatory variables. The marginal means were calculated intercepting the water treatment, while the significant estimates were obtained contrasting the differences form water and drought plots (Drought minus Control)). Significant p values are highlighted in bold. Abbreviations: Emmean = Estimated marginal means, SE = Standard errors, *P* = P. values, Amb = Ambient temperature; +2 °C = Ambient +2 °C warming; +HW = Ambient + heatwave warming (+10 °C); +2 °C+HW = Ambient +2 °C + heatwave warming; 1-month recovery = One month after drought; 4-months recovery = Four-months after drought.

| Season | Drought Treatment | Warming regimes | emmean | SE | *P* |
| --- | --- | --- | --- | --- | --- |
| Just before the drought | Control | Amb | -4.656 | 2.923 | 0.153 |
|  | Drought | Amb | -3.898 | 2.923 |  |
|  | Control | +2°C | -4.099 | 2.923 | 0.136 |
|  | Drought | +2°C | -4.889 | 2.923 |  |
|  | Control | +HW | -4.942 | 2.927 | 0.489 |
|  | Drought | +HW | -4.534 | 2.929 |  |
|  | Control | +2°C+HW | -4.488 | 2.923 | 0.517 |
|  | Drought | +2°C+HW | -4.144 | 2.923 |  |
| 1-Month recovery | **Control** | **Amb** | **-4.046** | **2.851** | **0.007** |
|  | **Drought** | **Amb** | **-2.966** | **2.851** |  |
|  | Control | +2°C | -3.245 | 2.851 | 0.575 |
|  | Drought | +2°C | -3.470 | 2.851 |  |
|  | Control | +HW | -3.542 | 2.845 | 0.109 |
|  | Drought | +HW | -4.199 | 2.851 |  |
|  | Control | +2°C+HW | -3.464 | 2.851 | 0.988 |
|  | Drought | +2°C+HW | -3.470 | 2.851 |  |
| 4-Months recovery | Control | Amb | -4.715 | 2.851 | 0.927 |
|  | Drought | Amb | -4.678 | 2.851 |  |
|  | **Control** | **+2°C** | **-3.779** | **2.851** | **0.028** |
|  | **Drought** | **+2°C** | **-4.658** | **2.851** |  |
|  | Control | +HW | -4.178 | 2.851 | 0.514 |
|  | Drought | +HW | -3.916 | 2.851 |  |
|  | **Control** | **+2°C+HW** | **-3.720** | **2.851** | **0.029** |
|  | **Drought** | **+2°C+HW** | **-4.595** | **2.851** |  |

Table S6: Estimated marginal means of drought by temperature and seasons on Simpson’s diversity (q=2). Output of the linear mixed-effects models testing invertebrate total abundances to warming, watering and season. The model included water treatment, warming regimes, harvest time point as explanatory variables. The marginal means were calculated intercepting the water treatment, while the significant estimates were obtained contrasting the differences form water and drought plots (Drought minus Control). Significant p values are highlighted in bold. Abbreviations: Emmean = Estimated marginal means, SE = Standard errors, *P* = P. values, Amb = Ambient temperature; +2 °C = Ambient +2 °C warming; +HW = Ambient + heatwave warming (+10 °C); +2 °C+HW = Ambient +2 °C + heatwave warming; 1-month recovery = One month after drought; 4-months recovery = Four-months after drought.

| Season | Drought Treatment | Warming regimes | emmean | SE | *P* |
| --- | --- | --- | --- | --- | --- |
| Just before the drought | **Control** | **Amb** | **1.224** | **3.068** | **0.043** |
|  | **Drought** | **Amb** | **2.323** | **3.068** |  |
|  | Control | +2°C | 1.877 | 3.068 | 0.245 |
|  | Drought | +2°C | 1.246 | 3.068 |  |
|  | Control | +HW | 1.243 | 3.072 | 0.623 |
|  | Drought | +HW | 1.541 | 3.074 |  |
|  | Control | +2°C+HW | 1.485 | 3.068 | 0.528 |
|  | Drought | +2°C+HW | 1.828 | 3.068 |  |
| 1-Month recovery | **Control** | **Amb** | **1.715** | **2.992** | **0.013** |
|  | **Drought** | **Amb** | **2.718** | **2.992** |  |
|  | Control | +2°C | 2.486 | 2.992 | 0.587 |
|  | Drought | +2°C | 2.267 | 2.992 |  |
|  | **Control** | **+HW** | **2.477** | **2.986** | **0.017** |
|  | **Drought** | **+HW** | **1.497** | **2.992** |  |
|  | Control | +2°C+HW | 2.237 | 2.992 | 0.958 |
|  | Drought | +2°C+HW | 2.215 | 2.992 |  |
| 4-Months recovery | Control | Amb | 1.143 | 2.992 | 0.760 |
|  | Drought | Amb | 1.020 | 2.992 |  |
|  | **Control** | **+2°C** | **2.054** | **2.992** | **0.022** |
|  | **Drought** | **+2°C** | **1.134** | **2.992** |  |
|  | Control | +HW | 1.715 | 2.992 | 0.593 |
|  | Drought | +HW | 1.930 | 2.992 |  |
|  | **Control** | **+2°C+HW** | **2.129** | **2.992** | **0.045** |
|  | **Drought** | **+2°C+HW** | **1.324** | **2.992** |  |

Table S7: Estimated marginal means of drought by temperature and seasons on Hymenoptera abundances. Output of the linear mixed-effects models testing invertebrate total abundances to warming, watering and season. The model included water treatment, warming regimes, harvest time point as explanatory variables. The marginal means were calculated intercepting the water treatment, while the significant estimates were obtained contrasting the differences form water and drought plots (Drought minus Control).Results are averaged over the levels of: Year and they are given on the log (not the response) scale. Significant p values are highlighted in bold. Abbreviations: Emmean = Estimated marginal means, SE = Standard errors, *P* = P. values, Amb = Ambient temperature; +2 °C = Ambient +2 °C warming; +HW = Ambient + heatwave warming (+10 °C); +2 °C+HW = Ambient +2 °C + heatwave warming; 1-month recovery = One month after drought; 4-months recovery = Four-months after drought.

| Season | Drought Treatment | Warming regimes | emmean | SE | P |
| --- | --- | --- | --- | --- | --- |
| Just before the drought | Control | Amb | 1.647 | 0.388 | 0.508 |
|  | Drought | Amb | 1.995 | 0.372 |  |
|  | Control | +2°C | 1.349 | 0.426 | 0.539 |
|  | Drought | +2°C | 1.696 | 0.387 |  |
|  | **Control** | **+HW** | **-0.297** | **0.696** | **0.013** |
|  | **Drought** | **+HW** | **1.722** | **0.455** |  |
|  | Control | +2°C+HW | 1.962 | 0.420 | 0.123 |
|  | Drought | +2°C+HW | 2.785 | 0.355 |  |
| 1-Month recovery | **Control** | **Amb** | **3.154** | **0.260** | **0.007** |
|  | **Drought** | **Amb** | **2.134** | **0.275** |  |
|  | Control | +2°C | 2.194 | 0.286 | 0.847 |
|  | Drought | +2°C | 2.271 | 0.286 |  |
|  | Control | +HW | 2.356 | 0.277 | 0.608 |
|  | Drought | +HW | 2.561 | 0.287 |  |
|  | Control | +2°C+HW | 2.742 | 0.261 | 0.285 |
|  | Drought | +2°C+HW | 2.336 | 0.278 |  |
| 4-Months recovery | Control | Amb | -0.508 | 0.484 | 0.564 |
|  | Drought | Amb | -0.936 | 0.564 |  |
|  | Control | +2°C | -0.353 | 0.460 | 0.150 |
|  | Drought | +2°C | -1.623 | 0.754 |  |
|  | Control | +HW | -0.108 | 0.425 | 0.944 |
|  | Drought | +HW | -0.150 | 0.424 |  |
|  | Control | +2°C+HW | 0.077 | 0.401 | 0.303 |
|  | Drought | +2°C+HW | -0.572 | 0.492 |  |

Table S8: Estimated marginal means of drought by temperature and seasons on Stylommatophora abundances. Output of the linear mixed-effects models testing invertebrate total abundances to warming, watering and season. The model included water treatment, warming regimes, harvest time point as explanatory variables. The marginal means were calculated intercepting the water treatment, while the significant estimates were obtained contrasting the differences form water and drought plots (Drought minus Control).Results are averaged over the levels of: Year and they are given on the log (not the response) scale. Significant p values are highlighted in bold. Abbreviations: Emmean = Estimated marginal means, SE = Standard errors, *P* = P. values, Amb = Ambient temperature; +2 °C = Ambient +2 °C warming; +HW = Ambient + heatwave warming (+10 °C); +2 °C+HW = Ambient +2 °C + heatwave warming; 1-month recovery = One month after drought; 4-months recovery = Four-months after drought.

| Season | Drought Treatment | Warming regimes | emmean | SE | P |
| --- | --- | --- | --- | --- | --- |
| Just before the drought | Control | Amb | 1.656 | 0.272 | 0.969 |
|  | Drought | Amb | 1.642 | 0.266 |  |
|  | **Control** | **+2°C** | **1.164** | **0.279** | **0.029** |
|  | **Drought** | **+2°C** | **2.010** | **0.290** |  |
|  | Control | +HW | 1.583 | 0.294 | 0.847 |
|  | Drought | +HW | 1.504 | 0.304 |  |
|  | Control | +2°C+HW | 1.260 | 0.275 | 0.871 |
|  | Drought | +2°C+HW | 1.320 | 0.276 |  |
| 1-Month recovery | Control | Amb | 1.469 | 0.245 | 0.056 |
|  | Drought | Amb | 0.721 | 0.311 |  |
|  | Control | +2°C | 0.033 | 0.337 | 0.360 |
|  | Drought | +2°C | -0.413 | 0.362 |  |
|  | **Control** | **+HW** | **0.367** | **0.319** | **0.015** |
|  | **Drought** | **+HW** | **-0.949** | **0.442** |  |
|  | Control | +2°C+HW | -0.475 | 0.360 | 0.884 |
|  | Drought | +2°C+HW | -0.552 | 0.391 |  |
| 4-Months recovery | Control | Amb | 2.071 | 0.225 | 0.095 |
|  | Drought | Amb | 1.536 | 0.236 |  |
|  | Control | +2°C | 1.885 | 0.232 | 0.198 |
|  | Drought | +2°C | 1.461 | 0.243 |  |
|  | **Control** | **+HW** | **2.197** | **0.222** | **0.011** |
|  | **Drought** | **+HW** | **1.364** | **0.249** |  |
|  | Control | +2°C+HW | 1.752 | 0.234 | 0.143 |
|  | Drought | +2°C+HW | 2.222 | 0.225 |  |

Table S9: Estimated marginal means of drought by temperature and seasons on Hemiptera abundances. Output of the linear mixed-effects models testing invertebrate total abundances to warming, watering and season. The model included water treatment, warming regimes, harvest time point as explanatory variables. The marginal means were calculated intercepting the water treatment, while the significant estimates were obtained contrasting the differences form water and drought plots (Drought minus Control). Results are averaged over the levels of: Year, Significant p values are highlighted in bold. Abbreviations: Emmean = Estimated marginal means, SE = Standard errors, *P* = P. values, Amb = Ambient temperature; +2 °C = Ambient +2 °C warming; +HW = Ambient + heatwave warming (+10 °C); +2 °C+HW = Ambient +2 °C + heatwave warming; 1-month recovery = One month after drought; 4-months recovery = Four-months after drought.

| Season | Drought Treatment | Warming regimes | emmean | SE | P |
| --- | --- | --- | --- | --- | --- |
| Just before the drought | Control | Amb | 1.803 | 0.492 | 0.127 |
|  | Drought | Amb | 2.709 | 0.370 |  |
|  | Control | +2°C | 2.122 | 0.453 | 0.965 |
|  | Drought | +2°C | 2.152 | 0.536 |  |
|  | Control | +HW | 2.571 | 0.478 | 0.327 |
|  | Drought | +HW | 1.884 | 0.533 |  |
|  | Control | +2°C+HW | 2.759 | 0.409 | 0.466 |
|  | Drought | +2°C+HW | 2.321 | 0.463 |  |
| 1-Month recovery | Control | Amb | 2.279 | 0.299 | 0.829 |
|  | Drought | Amb | 2.364 | 0.280 |  |
|  | Control | +2°C | 2.621 | 0.258 | 0.563 |
|  | Drought | +2°C | 2.814 | 0.240 |  |
|  | Control | +HW | 2.292 | 0.284 | 0.461 |
|  | Drought | +HW | 2.565 | 0.267 |  |
|  | Control | +2°C+HW | 2.602 | 0.264 | 0.243 |
|  | Drought | +2°C+HW | 2.151 | 0.303 |  |
| 4-Months recovery | Control | Amb | 1.649 | 0.363 | 0.604 |
|  | Drought | Amb | 1.385 | 0.382 |  |
|  | Control | +2°C | 1.153 | 0.412 | 0.734 |
|  | Drought | +2°C | 0.951 | 0.453 |  |
|  | Control | +HW | 1.170 | 0.411 | 0.369 |
|  | Drought | +HW | 0.598 | 0.509 |  |
|  | Control | +2°C+HW | 0.902 | 0.459 | 0.798 |
|  | Drought | +2°C+HW | 1.061 | 0.446 |  |

Table S10: Estimated marginal means of drought by temperature and seasons on Coleoptera abundances. Output of the linear mixed-effects models testing invertebrate total abundances to warming, watering and season. The model included water treatment, warming regimes, harvest time point as explanatory variables. The marginal means were calculated intercepting the water treatment, while the significant estimates were obtained contrasting the differences form water and drought plots (Drought minus Control).Results are averaged over the levels of: Year and they are given on the log (not the response) scale. Significant p values are highlighted in bold. Abbreviations: Emmean = Estimated marginal means, SE = Standard errors, *P* = P. values, Amb = Ambient temperature; +2 °C = Ambient +2 °C warming; +HW = Ambient + heatwave warming (+10 °C); +2 °C+HW = Ambient +2 °C + heatwave warming; 1-month recovery = One month after drought; 4-months recovery = Four-months after drought.

| Season | Drought Treatment | Warming regimes | emmean | SE | P |
| --- | --- | --- | --- | --- | --- |
| Just before the drought | Control | Amb | -0.093 | 0.620 | 0.486 |
|  | Drought | Amb | 0.467 | 0.526 |  |
|  | Control | +2°C | -0.786 | 0.796 | 0.348 |
|  | Drought | +2°C | 0.130 | 0.578 |  |
|  | Control | +HW | -1.256 | 1.080 | 0.064 |
|  | Drought | +HW | 0.919 | 0.475 |  |
|  | Control | +2°C+HW | -0.093 | 0.620 | 0.489 |
|  | Drought | +2°C+HW | -0.786 | 0.796 |  |
| 1-Month recovery | Control | Amb | 0.962 | 0.318 | 0.708 |
|  | Drought | Amb | 0.790 | 0.330 |  |
|  | Control | +2°C | 0.718 | 0.334 | 0.254 |
|  | Drought | +2°C | 1.233 | 0.304 |  |
|  | Control | +HW | 1.423 | 0.312 | 0.090 |
|  | Drought | +HW | 0.639 | 0.341 |  |
|  | Control | +2°C+HW | 1.238 | 0.303 | 0.700 |
|  | Drought | +2°C+HW | 1.070 | 0.313 |  |
| 4-Months recovery | Control | Amb | 0.060 | 0.394 | 0.889 |
|  | Drought | Amb | 0.138 | 0.394 |  |
|  | Control | +2°C | 1.304 | 0.306 | 0.104 |
|  | Drought | +2°C | 0.550 | 0.346 |  |
|  | Control | +HW | 0.503 | 0.351 | 0.729 |
|  | Drought | +HW | 0.327 | 0.368 |  |
|  | Control | +2°C+HW | 0.284 | 0.375 | 0.406 |
|  | Drought | +2°C+HW | -0.193 | 0.435 |  |

Table S11: Estimated marginal means of drought by temperature and seasons on the less abundant orders pulled abundances. Output of the linear mixed-effects models testing invertebrate total abundances to warming, watering and season. The model included water treatment, warming regimes, harvest time point as explanatory variables. The marginal means were calculated intercepting the water treatment, while the significant estimates were obtained contrasting the differences form water and drought plots (Drought minus Control).Results are averaged over the levels of: Year and they are given on the log (not the response) scale. Significant p values are highlighted in bold. Abbreviations: Emmean = Estimated marginal means, SE = Standard errors, *P* = P. values, Amb = Ambient temperature; +2 °C = Ambient +2 °C warming; +HW = Ambient + heatwave warming (+10 °C); +2 °C+HW = Ambient +2 °C + heatwave warming; 1-month recovery = One month after drought; 4-months recovery = Four-months after drought.

| Season | Drought Treatment | Warming regimes | emmean | SE | P |
| --- | --- | --- | --- | --- | --- |
| Just before the drought | Control | Amb | 0.977 | 0.507 | 0.266 |
|  | Drought | Amb | 1.732 | 0.476 |  |
|  | Control | +2°C | 0.999 | 0.501 | 0.054 |
|  | Drought | +2°C | -0.518 | 0.616 |  |
|  | Control | +HW | -0.274 | 0.655 | 0.996 |
|  | Drought | +HW | -18.759 | 3806.847 |  |
|  | Control | +2°C+HW | 1.321 | 0.495 | 0.613 |
|  | Drought | +2°C+HW | 1.670 | 0.514 |  |
| 1-Month recovery | Control | Amb | 1.811 | 0.358 | 0.894 |
|  | Drought | Amb | 1.744 | 0.357 |  |
|  | Control | +2°C | 1.873 | 0.351 | 0.675 |
|  | Drought | +2°C | 2.078 | 0.342 |  |
|  | Control | +HW | 0.779 | 0.434 | 0.194 |
|  | Drought | +HW | 1.512 | 0.357 |  |
|  | Control | +2°C+HW | 1.838 | 0.355 | 0.688 |
|  | Drought | +2°C+HW | 2.036 | 0.343 |  |
| 4-Months recovery | Control | Amb | 0.616 | 0.397 | 0.067 |
|  | Drought | Amb | 1.584 | 0.353 |  |
|  | Control | +2°C | 1.920 | 0.343 | 0.410 |
|  | Drought | +2°C | 2.338 | 0.378 |  |
|  | Control | +HW | 1.315 | 0.369 | 0.624 |
|  | Drought | +HW | 1.056 | 0.376 |  |
|  | Control | +2°C+HW | 1.439 | 0.358 | 0.322 |
|  | Drought | +2°C+HW | 1.937 | 0.349 |  |

Table S12: Estimated marginal means of temperature by recovery period on order rank difference from drought and control. Output of the linear mixed-effects models testing invertebrate total abundances to warming, watering and season. The model included water treatment, warming regimes, harvest time point as explanatory variables. The marginal means were calculated on warming effect while the significant estimates were obtained contrasting the differences form warmed plots and control (Ambient) ones. Significant p values are highlighted in bold. Abbreviations: Emmean = Estimated marginal means, SE = Standard errors, *P* = P. values, Amb = Ambient temperature; +2 °C = Ambient +2 °C warming; +HW = Ambient + heatwave warming (+10 °C); +2 °C+HW = Ambient +2 °C + heatwave warming; 1-month recovery = One month after drought; 4-months recovery = Four-months after drought.

| Season | Warming regimes | emmean | SE | *P* |
| --- | --- | --- | --- | --- |
| 1-Month recovery | Amb | 0.232 | 0.011 |  |
|  | +2°C | 0.223 | 0.011 | 0.929 |
|  | +HW | 0.226 | 0.011 | 0.982 |
|  | **+2°C+HW** | **0.192** | **0.011** | **0.040** |
| 4-Months recovery | Amb | 0.247 | 0.011 |  |
|  | +2°C | 0.268 | 0.011 | 0.514 |
|  | +HW | 0.255 | 0.011 | 0.951 |
|  | +2°C+HW | 0.234 | 0.011 | 0.826 |

Table S13: Estimated marginal means of temperature by recovery period on order cumulated curve difference from drought and control. Output of the linear mixed-effects models testing invertebrate total abundances to warming, watering and season. The model included water treatment, warming regimes, harvest time point as explanatory variables. The marginal means were calculated on warming effect while the significant estimates were obtained contrasting the differences form warmed plots and control (Ambient) ones. Significant p values are highlighted in bold. Abbreviations: Emmean = Estimated marginal means, SE = Standard errors, *P* = P. values, Amb = Ambient temperature; +2 °C = Ambient +2 °C warming; +HW = Ambient + heatwave warming (+10 °C); +2 °C+HW = Ambient +2 °C + heatwave warming; 1-month recovery = One month after drought; 4-months recovery = Four-months after drought.

| Season | Warming regimes |  | response | SE | P |
| --- | --- | --- | --- | --- | --- |
| 1-Month recovery | Amb |  | 15.882 | 2.361 |  |
|  | +2°C |  | 10.170 | 1.512 | 0.146 |
|  | +HW |  | 12.068 | 1.892 | 0.581 |
|  | +2°C+HW |  | 12.978 | 1.929 | 0.772 |
| 4-Months recovery | Amb |  | 6.745 | 1.003 |  |
|  | +2°C |  | 9.300 | 1.382 | 0.421 |
|  | +HW |  | 4.402 | 0.654 | 0.177 |
|  | +2°C+HW |  | 8.477 | 1.260 | 0.698 |

Table S14: Marginal and conditional r squares obtained in the regression models. We display per every regressions model (indexes and orders) the explained variance. The r squared are obtained with the package MuMIn v1.48.11.

|  | Metric | Marginal r^2^ | Conditional r^2^ |
| --- | --- | --- | --- |
| Diversity indexes | Abundance | 0.561 | 0.561 |
|  | Richness | 0.427 | 0.491 |
|  | Shannon | 0.399 | 0.508 |
|  | Simpson | 0.379 | 0.466 |
| Orders | Hemiptera | 0.464 | 0.469 |
|  | Stylommatophora | 0.797 | 0.817 |
|  | Hymenoptera | 0.713 | 0.777 |
|  | Coleoptera | 0.340 | 0.340 |

Table S15: Regressions between invertebrates total abundances and total above ground plant biomass. We display the estimated trends of drought in relation of the plant biomass, we contrasted according by temperature treatments and seasons. The generalised mixed effect model included water treatment, warming regimes, harvest time point, total plants above ground biomass and the year of collection as explanatory variables. The regression were tested if different from zero (*P*) and if different from control and drought (*P. contrast*). Significant *p. values* are highlighted in bold. Abbreviations: Emmean = Estimated marginal means, SE = Standard errors, AGB: Above Ground Biomass, Amb = Ambient temperature; +2 °C = Ambient +2 °C warming; +HW = Ambient + heatwave warming (+10 °C); +2 °C+HW = Ambient +2 °C + heatwave warming; 1-month recovery = One month after drought; 4-months recovery = Four-months after drought.

| Season | Drought Treatment | Warming regimes | Emmean | SE | AGB trend | *P.* | *P.* contrast |
| --- | --- | --- | --- | --- | --- | --- | --- |
| Just before the drought | Control | Amb | 3.211 | 0.292 | -0.001 | 0.883 | 0.879 |
|  | Drought | Amb | 3.579 | 0.337 | 0.000 | 0.943 |  |
|  | Control | +2°C | 2.594 | 0.386 | -0.008 | 0.235 | 0.180 |
|  | Drought | +2°C | 3.309 | 0.316 | 0.002 | 0.534 |  |
|  | Control | +HW | 3.096 | 0.319 | -0.001 | 0.741 | 0.733 |
|  | Drought | +HW | 2.988 | 0.431 | -0.006 | 0.663 |  |
|  | Control | +2°C+HW | 3.467 | 0.297 | -0.003 | 0.676 | 0.627 |
|  | Drought | +2°C+HW | 3.582 | 0.299 | 0.002 | 0.774 |  |
| 1-Month recovery | Control | Amb | 3.699 | 0.225 | 0.002 | 0.279 | 0.156 |
|  | Drought | Amb | 3.244 | 0.208 | -0.002 | 0.350 |  |
|  | Control | +2°C | 3.511 | 0.219 | -0.003 | 0.069 | 0.368 |
|  | Drought | +2°C | 3.580 | 0.204 | -0.001 | 0.760 |  |
|  | Control | +HW | 3.135 | 0.216 | -0.002 | 0.092 | 0.161 |
|  | Drought | +HW | 3.482 | 0.198 | 0.002 | 0.435 |  |
|  | Control | +2°C+HW | 3.506 | 0.197 | 0.002 | 0.289 | 0.616 |
|  | Drought | +2°C+HW | 3.301 | 0.207 | 0.000 | 0.972 |  |
| 4-Months recovery | Control | Amb | 2.890 | 0.197 | 0.004 | 0.089 | 0.908 |
|  | **Drought** | **Amb** | **2.685** | **0.199** | **0.004** | **0.033** |  |
|  | Control | +2°C | 2.895 | 0.226 | 0.006 | 0.100 | 0.651 |
|  | Drought | +2°C | 2.759 | 0.204 | 0.004 | 0.193 |  |
|  | **Control** | **+HW** | **2.998** | **0.204** | **0.011** | **<0.001** | 0.154 |
|  | **Drought** | **+HW** | **2.593** | **0.235** | **0.005** | **0.039** |  |
|  | **Control** | **+2°C+HW** | **2.880** | **0.202** | **0.011** | **<0.001** | 0.169 |
|  | **Drought** | **+2°C+HW** | **3.047** | **0.230** | **0.006** | **0.028** |  |

Table S16: Regressions between invertebrates orders richness and total above ground plant biomass. We display the estimated trends of drought in relation of the plant biomass, we contrasted according by temperature treatments and seasons. The generalised mixed effect model included water treatment, warming regimes, harvest time point, total plants above ground biomass and the year of collection as explanatory variables. The regression were tested if different from zero (*P*) and if different from control and drought (*P. contrast*). Significant *p. values* are highlighted in bold. Abbreviations: Emmean = Estimated marginal means, SE = Standard errors, AGB: Above Ground Biomass Amb = Ambient temperature; +2 °C = Ambient +2 °C warming; +HW = Ambient + heatwave warming (+10 °C); +2 °C+HW = Ambient +2 °C + heatwave warming; 1-month recovery = One month after drought; 4-months recovery = Four-months after drought.

| Season | Drought Treatment | Warming regimes | emmean | SE | AGB trend | *P.* | *P.* contrast |
| --- | --- | --- | --- | --- | --- | --- | --- |
| Just before the drought | Control | Amb | 3.483 | 0.614 | -0.013 | 0.246 | 0.195 |
|  | Drought | Amb | 4.099 | 0.707 | 0.008 | 0.487 |  |
|  | Control | +2°C | 2.577 | 0.813 | -0.029 | 0.054 | 0.088 |
|  | Drought | +2°C | 2.848 | 0.664 | -0.000 | 0.969 |  |
|  | Control | +HW | 2.497 | 0.670 | 0.007 | 0.382 | 0.135 |
|  | Drought | +HW | 2.019 | 0.900 | -0.039 | 0.187 |  |
|  | **Control** | **+2°C+HW** | **4.216** | **0.625** | **-0.028** | **0.033** | **0.030** |
|  | Drought | +2°C+HW | 3.639 | 0.628 | 0.017 | 0.296 |  |
| 1-Month recovery | **Control** | **Amb** | **4.878** | **0.491** | **0.011** | **0.019** | **0.011** |
|  | Drought | Amb | 5.875 | 0.459 | -0.006 | 0.204 |  |
|  | Control | +2°C | 6.140 | 0.480 | -0.002 | 0.478 | 0.511 |
|  | Drought | +2°C | 5.471 | 0.451 | 0.001 | 0.785 |  |
|  | Control | +HW | 4.468 | 0.473 | 0.001 | 0.779 | 0.677 |
|  | Drought | +HW | 5.031 | 0.441 | 0.003 | 0.557 |  |
|  | Control | +2°C+HW | 5.782 | 0.438 | 0.004 | 0.394 | 0.993 |
|  | Drought | +2°C+HW | 5.735 | 0.458 | 0.004 | 0.635 |  |
| 4-Months recovery | Control | Amb | 3.593 | 0.438 | -0.003 | 0.509 | 0.363 |
|  | Drought | Amb | 3.939 | 0.442 | 0.002 | 0.553 |  |
|  | **Control** | **+2°C** | **5.247** | **0.494** | **0.018** | **0.017** | 0.068 |
|  | Drought | +2°C | 3.909 | 0.452 | 0.001 | 0.935 |  |
|  | Control | +HW | 4.535 | 0.452 | 0.008 | 0.227 | 0.483 |
|  | Drought | +HW | 4.512 | 0.511 | 0.002 | 0.670 |  |
|  | Control | +2°C+HW | 5.071 | 0.447 | 0.004 | 0.444 | 0.745 |
|  | Drought | +2°C+HW | 3.685 | 0.498 | 0.002 | 0.719 |  |

Table S17: Regressions between invertebrates Shannon’s diversity (q=1) and total above ground plant biomass. We display the estimated trends of drought in relation of the plant biomass, we contrasted according by temperature treatments and seasons. The generalised mixed effect model included water treatment, warming regimes, harvest time point, total plants above ground biomass and the year of collection as explanatory variables. The regression were tested if different from zero (*P*) and if different from control and drought (*P. contrast*). Emmean = Estimated marginal means, SE = Standard errors, AGB: Above Ground Biomass Amb = Ambient temperature; +2 °C = Ambient +2 °C warming; +HW = Ambient + heatwave warming (+10 °C); +2 °C+HW = Ambient +2 °C + heatwave warming; 1-month recovery = One month after drought; 4-months recovery = Four-months after drought.

| Season | Drought Treatment | Warming regimes | emmean | SE | AGB trend | *P.* | *P.* contrast |
| --- | --- | --- | --- | --- | --- | --- | --- |
| Just before the drought | Control | Amb | 2.335 | 0.408 | -0.005 | 0.495 | 0.413 |
|  | Drought | Amb | 3.367 | 0.467 | 0.004 | 0.630 |  |
|  | Control | +2°C | 2.493 | 0.535 | -0.014 | 0.137 | 0.252 |
|  | Drought | +2°C | 2.206 | 0.440 | -0.002 | 0.684 |  |
|  | Control | +HW | 2.104 | 0.443 | 0.002 | 0.611 | 0.080 |
|  | Drought | +HW | 1.889 | 0.591 | -0.032 | 0.093 |  |
|  | Control | +2°C+HW | 2.769 | 0.415 | -0.014 | 0.109 | 0.051 |
|  | Drought | +2°C+HW | 2.825 | 0.417 | 0.012 | 0.235 |  |
| 1-Month recovery | Control | Amb | 2.926 | 0.333 | 0.004 | 0.209 | 0.314 |
|  | Drought | Amb | 4.340 | 0.313 | -0.000 | 0.867 |  |
|  | Control | +2°C | 4.048 | 0.326 | -0.001 | 0.772 | 0.376 |
|  | Drought | +2°C | 3.694 | 0.308 | 0.002 | 0.365 |  |
|  | Control | +HW | 3.591 | 0.322 | 0.002 | 0.254 | 0.774 |
|  | Drought | +HW | 2.943 | 0.302 | 0.001 | 0.826 |  |
|  | Control | +2°C+HW | 3.767 | 0.301 | -0.001 | 0.837 | 0.169 |
|  | Drought | +2°C+HW | 3.628 | 0.313 | 0.007 | 0.140 |  |
| 4-Months recovery | Control | Amb | 2.343 | 0.300 | -0.005 | 0.108 | 0.274 |
|  | Drought | Amb | 2.379 | 0.303 | -0.001 | 0.690 |  |
|  | **Control** | **+2°C** | **3.899** | **0.335** | **0.016** | **<0.001** | **0.002** |
|  | Drought | +2°C | 2.367 | 0.309 | -0.003 | 0.490 |  |
|  | Control | +HW | 2.858 | 0.309 | -0.006 | 0.162 | 0.354 |
|  | Drought | +HW | 3.201 | 0.345 | -0.001 | 0.755 |  |
|  | **Control** | **+2°C+HW** | **3.331** | **0.306** | **-0.009** | **0.015** | 0.069 |
|  | Drought | +2°C+HW | 2.490 | 0.336 | 4.28e-05 | 0.990 |  |

Table S18 Regressions between invertebrates Simpsons diversity (q=2) and total above ground plant biomass We display the estimated trends of drought in relation of the plant biomass, we contrasted according by temperature treatments and seasons. The generalised mixed effect model included water treatment, warming regimes, harvest time point, total plants above ground biomass and the year of collection as explanatory variables. The regression were tested if different from zero (*P*) and if different from control and drought (*P. contrast*). Significant *p. values* are highlighted in bold. Abbreviations: Emmean = Estimated marginal means, SE = Standard errors, AGB: Above Ground Biomass Amb = Ambient temperature; +2 °C = Ambient +2 °C warming; +HW = Ambient + heatwave warming (+10 °C); +2 °C+HW = Ambient +2 °C + heatwave warming; 1-month recovery = One month after drought; 4-months recovery = Four-months after drought.

| Season | Drought Treatment | Warming regimes | emmean | SE | AGB trend | *P.* | *P.* contrast |
| --- | --- | --- | --- | --- | --- | --- | --- |
| Just before the drought | Control | Amb | 0.636 | 0.145 | -0.001 | 0.606 | 0.699 |
|  | Drought | Amb | 1.088 | 0.167 | 0.000 | 0.961 |  |
|  | Control | +2°C | 0.781 | 0.191 | -0.003 | 0.318 | 0.489 |
|  | Drought | +2°C | 0.691 | 0.157 | -0.001 | 0.651 |  |
|  | Control | +HW | 0.669 | 0.158 | 0.001 | 0.686 | 0.060 |
|  | Drought | +HW | 0.523 | 0.211 | -0.013 | 0.067 |  |
|  | Control | +2°C+HW | 0.797 | 0.147 | -0.003 | 0.311 | 0.125 |
|  | Drought | +2°C+HW | 0.839 | 0.148 | 0.004 | 0.258 |  |
| 1-Month recovery | Control | Amb | 0.817 | 0.117 | 0.001 | 0.409 | 0.809 |
|  | Drought | Amb | 1.238 | 0.109 | 0.001 | 0.623 |  |
|  | Control | +2°C | 1.169 | 0.114 | -0.000 | 0.884 | 0.508 |
|  | Drought | +2°C | 1.058 | 0.108 | 0.001 | 0.463 |  |
|  | Control | +HW | 1.129 | 0.113 | 0.001 | 0.262 | 0.934 |
|  | Drought | +HW | 0.782 | 0.105 | 0.001 | 0.538 |  |
|  | Control | +2°C+HW | 1.066 | 0.105 | -0.000 | 0.727 | 0.198 |
|  | Drought | +2°C+HW | 1.016 | 0.109 | 0.002 | 0.198 |  |
| 4-Months recovery | **Control** | **Amb** | **0.630** | **0.105** | **-0.002** | **0.048** | 0.072 |
|  | Drought | Amb | 0.590 | 0.106 | 0.000 | 0.803 |  |
|  | **Control** | **+2°C** | **1.138** | **0.118** | **0.005** | **0.007** | **0.008** |
|  | Drought | +2°C | 0.609 | 0.108 | -0.001 | 0.414 |  |
|  | **Control** | **+HW** | **0.806** | **0.108** | **-0.003** | **0.042** | 0.331 |
|  | Drought | +HW | 0.879 | 0.121 | -0.001 | 0.290 |  |
|  | **Control** | **+2°C+HW** | **0.952** | **0.107** | **-0.004** | **0.001** | **0.012** |
|  | Drought | +2°C+HW | 0.711 | 0.118 | 0.000 | 0.929 |  |

Table S19 Regressions between Hemiptera frequencies and total above ground plant biomass We display the estimated trends of drought in relation of the plant biomass, we contrasted according by temperature treatments and seasons. The generalised mixed effect model included water treatment, warming regimes, harvest time point, total plants above ground biomass and the year of collection as explanatory variables. The regression were tested if different from zero (*P*) and if different from control and drought (*P. contrast*). Significant *p. values* are highlighted in bold. Abbreviations: Emmean = Estimated marginal means, SE = Standard errors, AGB: Above Ground Biomass Amb = Ambient temperature; +2 °C = Ambient +2 °C warming; +HW = Ambient + heatwave warming (+10 °C); +2 °C+HW = Ambient +2 °C + heatwave warming; 1-month recovery = One month after drought; 4-months recovery = Four-months after drought.

| Season | Drought Treatment | Warming regimes | emmean | SE | AGB trend | *P.* | *P.* contrast |
| --- | --- | --- | --- | --- | --- | --- | --- |
| Just before the drought | Control | Amb | -0.085 | 0.343 | 0.004 | 0.564 | 0.232 |
|  | Drought | Amb | 0.514 | 0.397 | -0.007 | 0.272 |  |
|  | Control | +2°C | 0.219 | 0.454 | 0.001 | 0.915 | 0.774 |
|  | Drought | +2°C | 0.409 | 0.372 | 0.004 | 0.401 |  |
|  | Control | +HW | 0.632 | 0.375 | -0.002 | 0.687 | 0.060 |
|  | **Drought** | **+HW** | **-0.359** | **0.510** | **-0.034** | **0.041** |  |
|  | Control | +2°C+HW | 0.781 | 0.349 | 0.011 | 0.128 | 0.072 |
|  | Drought | +2°C+HW | 0.651 | 0.352 | -0.010 | 0.289 |  |
| 1-Month recovery | **Control** | **Amb** | **-0.024** | **0.265** | **0.006** | **0.024** | 0.293 |
|  | Drought | Amb | 0.336 | 0.246 | 0.002 | 0.449 |  |
|  | Control | +2°C | 0.825 | 0.259 | -0.002 | 0.212 | 0.957 |
|  | Drought | +2°C | 0.870 | 0.241 | -0.002 | 0.357 |  |
|  | Control | +HW | 0.215 | 0.254 | -0.001 | 0.451 | **0.035** |
|  | **Drought** | **+HW** | **0.778** | **0.234** | **0.006** | **0.049** |  |
|  | Control | +2°C+HW | 0.633 | 0.233 | 0.005 | 0.077 | 0.711 |
|  | Drought | +2°C+HW | 0.074 | 0.245 | 0.003 | 0.555 |  |
| 4-Months recovery | Control | Amb | -0.295 | 0.232 | -0.002 | 0.370 | 0.948 |
|  | Drought | Amb | -0.483 | 0.235 | -0.002 | 0.265 |  |
|  | Control | +2°C | -0.529 | 0.267 | 0.005 | 0.243 | 0.155 |
|  | Drought | +2°C | -0.778 | 0.242 | -0.003 | 0.424 |  |
|  | Control | +HW | -0.608 | 0.241 | 0.004 | 0.348 | 0.792 |
|  | Drought | +HW | -0.830 | 0.278 | 0.002 | 0.442 |  |
|  | **Control** | **+2°C+HW** | **-0.524** | **0.238** | **0.010** | **0.003** | 0.127 |
|  | Drought | +2°C+HW | -0.526 | 0.271 | 0.003 | 0.326 |  |

Table S20 Regressions between Stylommatophora frequencies and total above ground plant biomass We display the estimated trends of drought in relation of the plant biomass, we contrasted according by temperature treatments and seasons. The generalised mixed effect model included water treatment, warming regimes, harvest time point, total plants above ground biomass and the year of collection as explanatory variables. The regression were tested if different from zero (*P*) and if different from control and drought (*P. contrast*). Significant *p. values* are highlighted in bold. Abbreviations: Emmean = Estimated marginal means, SE = Standard errors, AGB: Above Ground Biomass Amb = Ambient temperature; +2 °C = Ambient +2 °C warming; +HW = Ambient + heatwave warming (+10 °C); +2 °C+HW = Ambient +2 °C + heatwave warming; 1-month recovery = One month after drought; 4-months recovery = Four-months after drought.

| Season | Drought Treatment | Warming regimes | emmean | SE | AGB trend | *P.* | *P.* contrast |
| --- | --- | --- | --- | --- | --- | --- | --- |
| Just before the drought | **Control** | **Amb** | **0.310** | **0.211** | **0.009** | **0.028** | **0.047** |
|  | Drought | Amb | 0.373 | 0.243 | -0.003 | 0.526 |  |
|  | Control | +2°C | -0.286 | 0.278 | -0.009 | 0.068 | 0.179 |
|  | Drought | +2°C | 0.232 | 0.228 | -0.002 | 0.542 |  |
|  | Control | +HW | 0.442 | 0.230 | -0.001 | 0.807 | 0.388 |
|  | Drought | +HW | 0.494 | 0.310 | 0.008 | 0.407 |  |
|  | Control | +2°C+HW | 0.151 | 0.214 | 0.001 | 0.837 | 0.429 |
|  | Drought | +2°C+HW | 0.076 | 0.216 | 0.006 | 0.235 |  |
| 1-Month recovery | Control | Amb | 0.102 | 0.168 | -0.001 | 0.732 | 0.248 |
|  | **Drought** | **Amb** | **-0.416** | **0.157** | **-0.003** | **0.044** |  |
|  | Control | +2°C | -0.790 | 0.164 | 0.001 | 0.309 | 0.163 |
|  | Drought | +2°C | -0.691 | 0.154 | -0.001 | 0.325 |  |
|  | Control | +HW | -0.497 | 0.162 | 0.001 | 0.319 | 0.204 |
|  | Drought | +HW | -1.058 | 0.150 | -0.002 | 0.349 |  |
|  | Control | +2°C+HW | -0.803 | 0.149 | 0.000 | 0.775 | 0.578 |
|  | Drought | +2°C+HW | -0.888 | 0.156 | 0.002 | 0.421 |  |
| 4-Months recovery | **Control** | **Amb** | **0.411** | **0.149** | **0.004** | **0.009** | 0.620 |
|  | **Drought** | **Amb** | **0.112** | **0.151** | **0.003** | **0.005** |  |
|  | **Control** | **+2°C** | 0.113 | 0.169 | -0.003 | 0.330 | 0.056 |
|  | Drought | +2°C | 0.069 | 0.154 | 0.004 | 0.074 |  |
|  | **Control** | **+HW** | **0.617** | **0.154** | **0.006** | **0.008** | 0.167 |
|  | Drought | +HW | 0.116 | 0.175 | 0.002 | 0.228 |  |
|  | **Control** | **+2°C+HW** | **0.308** | **0.153** | **0.005** | **0.018** | 0.485 |
|  | Drought | +2°C+HW | 0.592 | 0.170 | 0.003 | 0.118 |  |

Table S21 Regressions between Hymenoptera frequencies and total above ground plant biomass We display the estimated trends of drought in relation of the plant biomass, we contrasted according by temperature treatments and seasons. The generalised mixed effect model included water treatment, warming regimes, harvest time point, total plants above ground biomass and the year of collection as explanatory variables. The regression were tested if different from zero (*P*) and if different from control and drought (*P. contrast*). Significant *p. values* are highlighted in bold. Abbreviations: Emmean = Estimated marginal means, SE = Standard errors, AGB: Above Ground Biomass Amb = Ambient temperature; +2 °C = Ambient +2 °C warming; +HW = Ambient + heatwave warming (+10 °C); +2 °C+HW = Ambient +2 °C + heatwave warming; 1-month recovery = One month after drought; 4-months recovery = Four-months after drought.

| Season | Drought Treatment | Warming regimes | emmean | SE | AGB trend | *P.* | *P.* contrast |
| --- | --- | --- | --- | --- | --- | --- | --- |
| Just before the drought | Control | Amb | 0.345 | 0.252 | 0.000 | 0.981 | 0.232 |
|  | Drought | Amb | 0.546 | 0.287 | -0.002 | 0.691 |  |
|  | Control | +2°C | -0.252 | 0.326 | -0.006 | 0.336 | 0.774 |
|  | Drought | +2°C | 0.331 | 0.272 | -0.002 | 0.551 |  |
|  | Control | +HW | -0.768 | 0.273 | 0.003 | 0.257 | 0.060 |
|  | Drought | +HW | 0.162 | 0.362 | -0.009 | 0.460 |  |
|  | **Control** | **+2°C+HW** | **0.331** | **0.256** | **-0.014** | **0.005** | 0.072 |
|  | Drought | +2°C+HW | 1.013 | 0.257 | 0.007 | 0.238 |  |
| 1-Month recovery | Control | Amb | 1.235 | 0.208 | 0.002 | 0.200 | 0.293 |
|  | Drought | Amb | 0.492 | 0.196 | 0.001 | 0.591 |  |
|  | **Control** | **+2°C** | **0.794** | **0.204** | **-0.005** | **<0.001** | 0.957 |
|  | Drought | +2°C | 0.541 | 0.194 | -0.000 | 0.869 |  |
|  | **Control** | **+HW** | **0.805** | **0.202** | **-0.002** | **0.016** | **0.035** |
|  | Drought | +HW | 0.760 | 0.190 | 0.003 | 0.242 |  |
|  | Control | +2°C+HW | 1.085 | 0.189 | 0.003 | 0.086 | 0.711 |
|  | Drought | +2°C+HW | 0.723 | 0.196 | -0.001 | 0.720 |  |
| 4-Months recovery | Control | Amb | -0.887 | 0.189 | 0.002 | 0.202 | 0.948 |
|  | Drought | Amb | -0.939 | 0.191 | 0.001 | 0.374 |  |
|  | **Control** | **+2°C** | **-0.646** | **0.209** | **0.008** | **0.009** | 0.155 |
|  | Drought | +2°C | -1.035 | 0.194 | 0.003 | 0.313 |  |
|  | Control | +HW | -0.732 | 0.194 | 0.004 | 0.090 | 0.792 |
|  | Drought | +HW | -0.661 | 0.215 | 0.001 | 0.463 |  |
|  | Control | +2°C+HW | -0.666 | 0.192 | 0.004 | 0.085 | 0.127 |
|  | Drought | +2°C+HW | -0.896 | 0.210 | 0.002 | 0.441 |  |

Table S22 Regressions between Coleoptera frequencies and total above ground plant biomass We display the estimated trends of drought in relation of the plant biomass, we contrasted according by temperature treatments and seasons. The generalised mixed effect model included water treatment, warming regimes, harvest time point, total plants above ground biomass and the year of collection as explanatory variables. The regression were tested if different from zero (*P*) and if different from control and drought (*P. contrast*). Significant *p. values* are highlighted in bold. Abbreviations: Emmean = Estimated marginal means, SE = Standard errors, AGB: Above Ground Biomass Amb = Ambient temperature; +2 °C = Ambient +2 °C warming; +HW = Ambient + heatwave warming (+10 °C); +2 °C+HW = Ambient +2 °C + heatwave warming; 1-month recovery = One month after drought; 4-months recovery = Four-months after drought.

| Season | Drought Treatment | Warming regimes | emmean | SE | AGB trend | *P.* | *P.* contrast |
| --- | --- | --- | --- | --- | --- | --- | --- |
| Just before the drought | Control | Amb | -0.431 | 0.380 | -0.007 | 0.306 | 0.625 |
|  | Drought | Amb | -0.221 | 0.440 | -0.002 | 0.774 |  |
|  | Control | +2°C | -1.189 | 0.503 | -0.015 | 0.113 | 0.164 |
|  | Drought | +2°C | -0.257 | 0.413 | -0.000 | 0.962 |  |
|  | Control | +HW | -0.716 | 0.416 | -0.000 | 0.967 | **0.027** |
|  | **Drought** | **+HW** | **-0.590** | **0.563** | **-0.043** | **0.022** |  |
|  | Control | +2°C+HW | -0.119 | 0.387 | -0.011 | 0.187 | 0.186 |
|  | Drought | +2°C+HW | -0.725 | 0.390 | 0.006 | 0.526 |  |
| 1-Month recovery | Control | Amb | 0.402 | 0.293 | -0.003 | 0.261 | 0.393 |
|  | Drought | Amb | 0.362 | 0.271 | 0.000 | 0.927 |  |
|  | Control | +2°C | 0.274 | 0.286 | -0.001 | 0.785 | 0.663 |
|  | Drought | +2°C | 0.625 | 0.266 | 0.001 | 0.734 |  |
|  | Control | +HW | 0.690 | 0.281 | -0.002 | 0.345 | 0.359 |
|  | Drought | +HW | 0.182 | 0.259 | 0.002 | 0.561 |  |
|  | Control | +2°C+HW | 0.666 | 0.257 | 0.002 | 0.504 | 0.911 |
|  | Drought | +2°C+HW | 0.580 | 0.270 | 0.001 | 0.797 |  |
| 4-Months recovery | Control | Amb | -0.372 | 0.257 | -0.003 | 0.331 | **0.018** |
|  | **Drought** | **Amb** | **-0.394** | **0.260** | **0.006** | **0.011** |  |
|  | Control | +2°C | 0.710 | 0.295 | 0.007 | 0.167 | 0.064 |
|  | Drought | +2°C | -0.256 | 0.267 | -0.005 | 0.231 |  |
|  | Control | +HW | -0.050 | 0.266 | -0.004 | 0.287 | 0.230 |
|  | Drought | +HW | -0.035 | 0.307 | 0.002 | 0.589 |  |
|  | Control | +2°C+HW | -0.220 | 0.263 | 0.001 | 0.837 | 0.248 |
|  | Drought | +2°C+HW | -0.346 | 0.300 | 0.006 | 0.059 |  |
